# Use of Substrate Analogues and X-ray spectroscopy Reveals an all Ferrous C-Cluster in CO Dehydrogenase

**DOI:** 10.64898/2026.01.21.700957

**Authors:** Macon Abernathy, Kareem Aboulhosn, Christopher Ohmer, Stephen Ragsdale, Ritimukta Sarangi

## Abstract

Carbon monoxide (CO) dehydrogenase (CODH) plays a key role in prokaryotic one-carbon metabolism by detoxifying CO and by driving CO_2_ reduction coupled to ATP production in the Wood-Ljungdahl Pathway. Here we focus on a Ni-Fe CODH (CODH-II), with an active site C-cluster, which is a [NiFe_4_S_4_] cluster arranged as a [NiFe_3_S_4_] subcluster with an additional, unique pendant Fe, e.g., [Fe_3_S_4_-Fe_u_]. It catalyzes the reversible reduction of CO_2_ to CO without the requirement for an overpotential and with insignificant proton reduction. The redox states associated with catalysis are defined as C_red1_ and C_red2_. Despite crystal structures with near 1.0 A resolution, it has been a long-standing question where the electrons in these catalytically relevant redox states are stored within the C cluster. Using X-ray absorption spectroscopy (XAS), EPR, and substrate-analogue binding measurements, we clarify the electronic structure of these catalytically active states in addition to the resting state of CODH. We rule out recent postulates that catalysis involves a Ni^0^ state, a metal-metal bond, or a hydride intermediate. We reveal that CODH rests in the diamagnetic C_ox_ form, which contains Ni^2+^ and an oxidized [Fe_3_S_4_-Fe_u_]^2+^ cluster. Then, the C-cluster undergoes reductive activation on Fe to form paramagnetic Cred1, which binds CO and analog cyanide. Generation of C_red2_, which binds CO_2_ and its analog cyanate, involves two sequential valence-localized electron transfers, generating Ni^1+^ and then [Fe_3_S_4_-Fe_u_], forming an all-ferrous cluster. Our work sheds light on how CODH avoids the thermodynamically unfavorable generation of a CO_2_ radical anion intermediate formed in other catalytic systems by stabilizing electron density in the heterometallic C-cluster. We also highlight the importance of high-resolution XAS and use of substrate analogs to reveal the sequential, valence-localized electron transfers that occur during redox-dependent CODH catalysis.

**TOC Figure:** 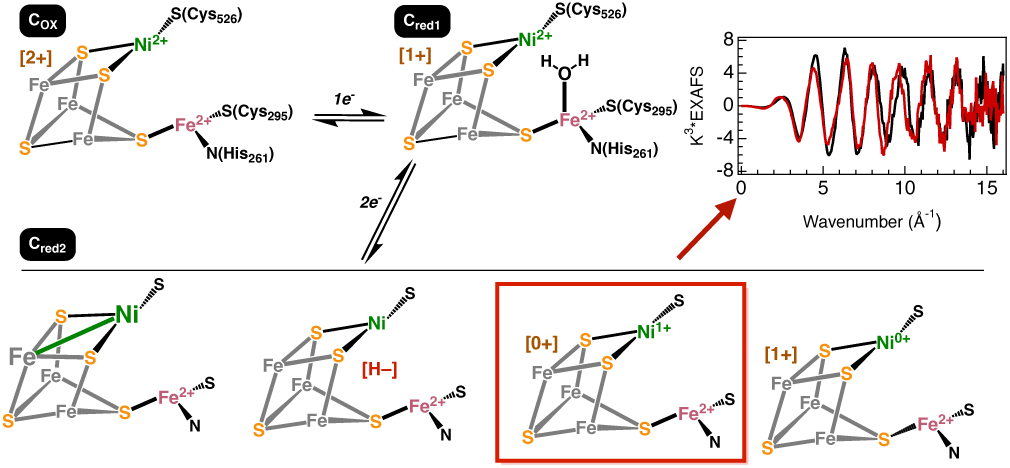

## Introduction

Carbon Monoxide Dehydrogenase (CODH) is a homodimeric enzyme which uses a unique [NiFe_3_S_4_-Fe_u_] containing C cluster to reversibly reduce CO_2_ to CO with negligible overpotential, thus catalyzing the biological equivalent of the industrial water-gas shift reaction. CODH couples to several important pathways. In the Wood-Ljungdahl pathway (WLP), CODH forms a complex with Acetyl CoA-Synthase (ACS), which captures the CO from an intersubunit CODH/ACS channel and catalyzes its condensation with a methyl group and CoA to form acetyl CoA. The WLP is the predominant mode of anaerobic carbon fixation, responsible for producing more than 10^13^ kg yr^-1^ of acetic acid globally. ^1,2^ CODH, ACS and the other enzymes composing the WLP are ancient, tracing their lineage directly to the last universal common ancestor, and are found today in both the bacterial and archaeal domains. ^3,4^ There is substantial industrial interest in the pathway as an efficient means to produce short chain carbon feedstocks for manufacturing or energy generation. ^5–8^ CODH also exists as free-standing enzyme (like the CODH-II from *Carboxydothermus hydrogenoformans* studied here) that couples to hydrogenase to catalyze proton reduction to H_2_.

In all its manifestations, the CODH dimer contains three [Fe_4_S_4_] clusters: the D cluster sits at the interface of the two monomeric units and facilitates electron transfer to and from external mediators to each of two B clusters, one per monomer. Each B cluster in turn shuttles electron to its respective C cluster – the site of CO_2_ reduction. We provide a detailed structural summary of these clusters in the supplementary information. The consensus structure of the C cluster is a [NiFe_3_S_4_-Fe^2+^] distorted cubane, in which the Ni is incorporated into the cubane and trigonally bound to two µ^3^-sulfides and C526 (PDB 4UDX). An additional ferrous iron (Fe_u_) is linked to the cluster via a single µ^3^-sulfide. ^9,10^ The C-cluster has been the focus of intense structural, electrochemical and spectroscopic studies. ^11–16^ These studies have greatly advanced our understanding of the structure of CODH, and potential binding modes for substrates and substrate mimics, and are summarized in the supplementary information. The C cluster can adopt one of four redox states, each corresponding to a stage in its catalytic cycle (Figure 1), exhibiting distinct spectroscopic signatures and redox potentials. In *Carboxydothermus hydrogenoformans*, the inactive, oxidized form of the enzyme (C_ox_) is reduced by one electron to the paramagnetic species C_red1_ (E°’ ∼ –100 mV), ^17^ which is responsible for binding CO. Further reduction by two electrons yields another paramagnetic species C_red2_ (E°’ ∼ –530 mV), ^18,19^ which binds CO_2_. A diamagnetic transient species, C_int_ has been proposed to form by the one electron reduction of C_red1_. A literature review of the physical properties of CODH is presented in SI Section II.

**Figure 1.**
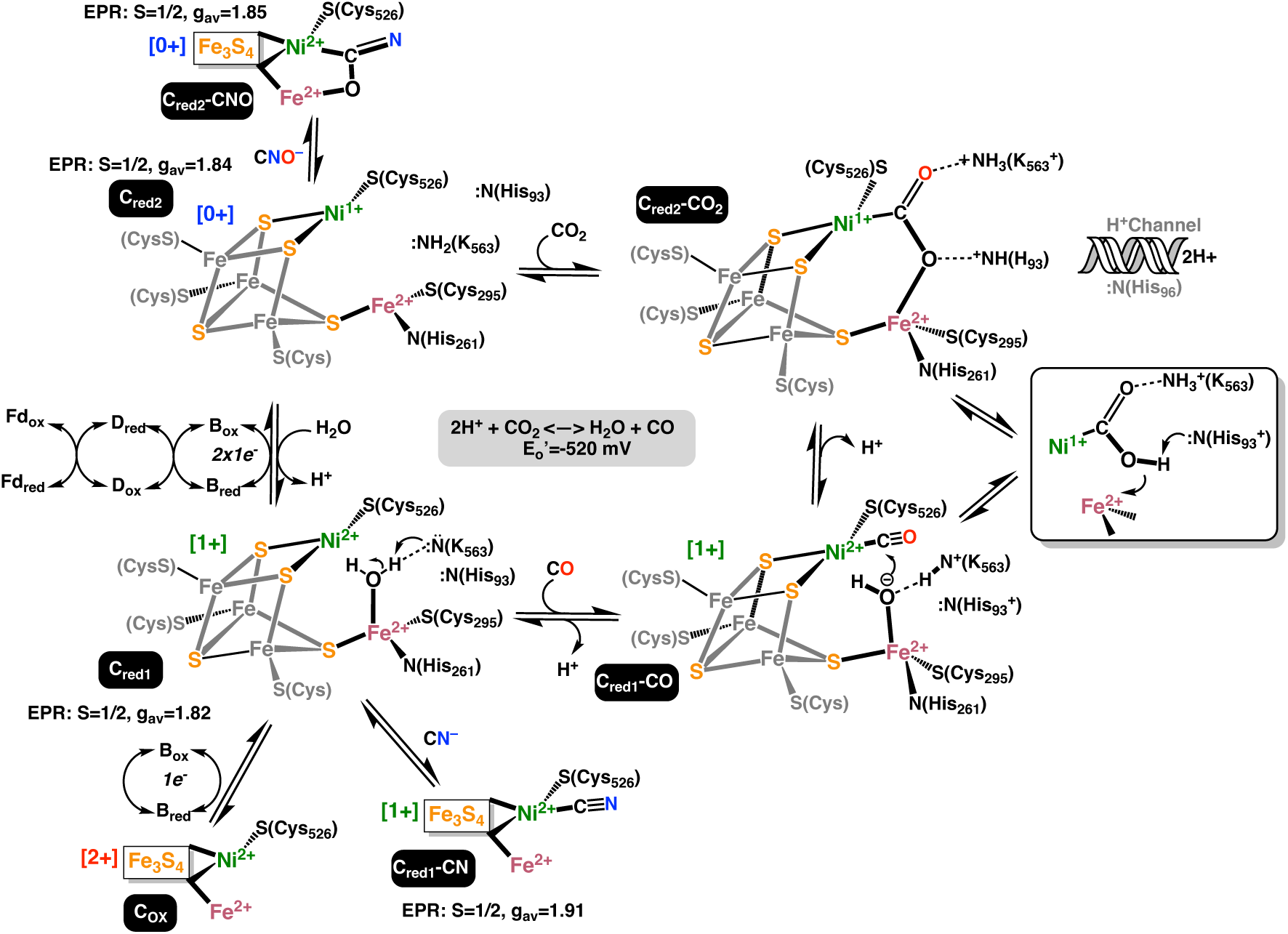
The mechanism of CODH highlighting the major catalytic intermediates of C cluster with oxidation state assignments from this work.

However, despite X-ray diffraction (XRD) structures at 1.0-1.3 Å overall resolution, ^20^ there remain long-standing questions as to where the electrons in the catalytically relevant redox states (C_red1_ and C_red2_) are stored within the C-cluster. Current postulates suggest that the electronic structure of Cred2 includes Ni^0^, ^21^ a metal-metal bond, ^22^ or a bound hydride. ^23^ The work described here provides evidence ruling out these hypotheses and offers an alternative. They have also sparked debate over the structure of the active site, including the binding mode of CO/CN^−^ and the presence of a bridging S^2-^ or Cl^-^ between the Ni and Fe_u_. ^12,15,16,24,25^ However, even the crystal structures with the best resolution suffer from significant uncertainty in the structure of the C cluster, exhibiting multiple conformations for key atoms. ^11,13,15,24,26^ Different structures have been obtained from similar treatments, for example poising at -320 mV with DTT to recreate the C_red1_ intermediate has resulted in structures with and without a bridging S^2-^ or Cl^-^ between the Ni and Fe_u_. ^11,24^ A recent crystallographic study has proposed several new intermediate states in the catalytic mechanism. ^20^ But this too suffers from inconsistencies with prior data, for example a C_red1_ state (PDB: 9FPG) reduced with Ti(III) at pH 6.0 yielded a bridging -OH ligand between the Ni and Fe_u_, inconsistent with prior C_red1_ structures (PDB: 3B53, 1SU7). Furthermore, reported structures from that study such as PDB: 9FPO exhibit internal inconsistencies in the modelling of a C cluster simultaneously in a CO and CO_2_ bound state yielding unlikely bond distances, which are addressed here. A systematic assessment of the diversity of crystallographic distances for all CODH structures in the PDB underscores this discrepancy from available crystallographic data on all CODH structures. The C cluster region, when parsed into any alternate location configurations specified in the PDB file reveals wide discrepancy in Ni-S and Ni-Fe distances Figure 2 (Table S1, Figure S1, SI Section I), with standard deviations of ∼0.1 Å for Ni-S and ∼0.25 Å and ∼0.22 Å for Fe_u_ and Fe_2_. We note that studies presenting new structures rarely provide explanations for disagreements with previously reported structures for the same species and state. No doubt that the spread in observed interatomic distances reflects differences in crystallization conditions, resolution, fit quality, redox poise, photodamage and species of origin. For the biochemist concerned with relating the structure of CODH with its function in the solution phase, this poses additional challenges in the development of the underlying catalytic mechanism.

**Figure 2.**
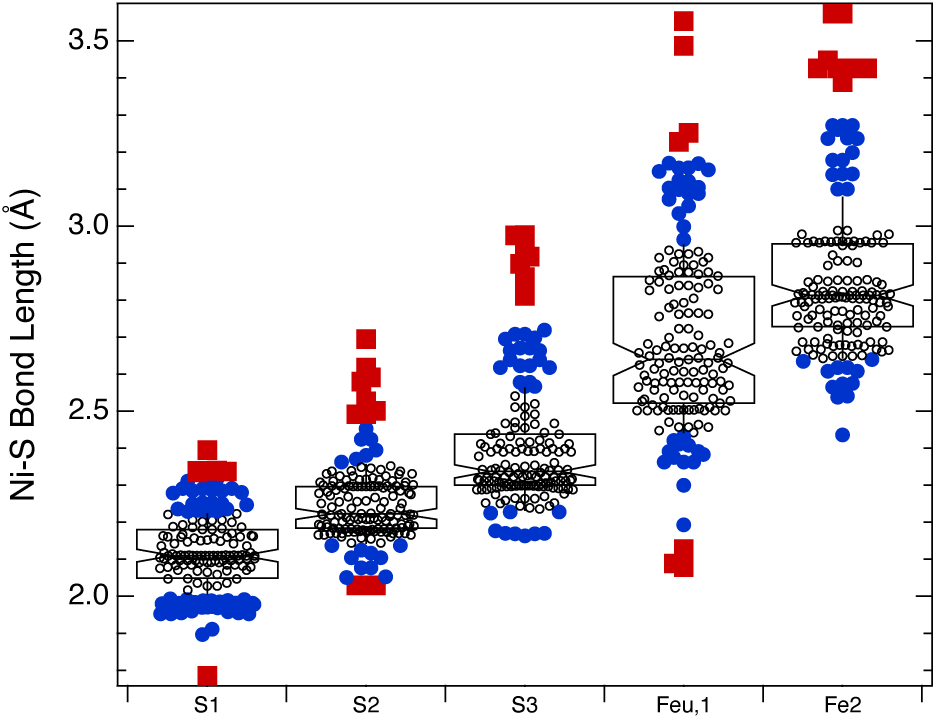
A survey of Ni-S/Fe distances from all available PDB structures of the C cluster. Numbering corresponds to Schematic S1. The central box contains markings for the 1st and 3rd quartile, and median value. Shaded circles represent values more than one SD away from the mean, and shaded squares represent values more than 2 SD from the mean. The notches at the median value represent the 95% confidence interval for the median value.

Spectroscopic investigations using Ni K-edge XAS have not necessarily fared better, with existing data often showing little to no change among the various intermediates, suggesting a lack of conversion. ^27–30^ Thus, there is a strong need for a detailed structural description of the C cluster, particularly at the catalytic Ni in its substrate free states. In this study, we present a thorough EPR and XAS characterization of the C cluster in the C_ox_, C_red1_ and C_red2_ states and in the presence of the inhibitory analogs for CO (CN^−^) and CO_2_ (CNO^−^) in solution. The structural and redox assignments described here are supported by previous rigorous electrochemical analyses, ^16^ demystifying the significant incongruence in the crystallography literature, and laying a solid foundation for developing our understanding of the underlying mechanism.

## 1. Materials and Methods

CODH-II was expressed and purified from *C. hydrogenoformans* Z-2901 using previously established protocols. ^17^ The CODH-C_ox_ state is prepared by incubating 2 mM CODH-II with 10 mM resazurin (E^0’^ = 110 mV) for 15 minutes and then freezing in LN2. The CODH-C_red1_ state is prepared by incubating 2 mM CODH-II with 20 mM dithiothreitol and 1 mM benzyl viologen (E^0’^ = 370 mV) for 15 minutes, then freezing in LN2. UV-Vis measurement confirmed that half of the benzyl viologen was poised in the reduced state under these conditions. The CODH-C_red2_ state is prepared by incubating 2 mM CODH-II with 20 mM dithionite and 1 mM triquat for 15 minutes then freezing in LN2 (E^0’^ = -540 mV). Protein concentration was determined using Rose-Bengal assay. Specific activity of CODH-II varied between 500-775 U/mg. The CODH-C_red1_-CN XAS sample is prepared by incubating 2 mM CODH-II with 20 mM dithiothreitol, 1 mM benzyl viologen, and 50 mM KCN for 15 minutes then freezing in LN2 in the anaerobic chamber. The CODH-C_red2_-OCN XAS sample is prepared by incubating 2 mM CODH-II with 20 mM dithionite, 1 mM triquat, and 250 mM KOCN for 15 minutes and then freezing in LN2. All XAS measurements were collected on beamline 7-3 at the Stanford Synchrotron Radiation Lightsource, and HERFD were collected at beamline 15-2. Pyspline, Athena and EXAFSPAK were used for data processing and analysis. ^69–71^ X-band EPR measurements were collected with a Bruker ECS106 spectrometer. Additional details on sample preparation and experimental procedures are provided in Section I of the Supplementary Information.

## 2. Redox Poising and EPR Spectroscopy Define the Paramagnetic States of CODH-II

The redox properties of the 3 paramagnetic centers in CODH have been previously investigated in orthologs. The B- and D-clusters have a midpoint redox potential ([Fe_4_S_4_]^2+/1+^) of -440 mV and -530 mV respectively, ^9,18,31^ although they vary between *Rhodospirillum rubrum* and *Neomoorella thermoacetica (f. Moorella thermoacetica)*. ^18,32,33^ Of the four putative redox states of the C-cluster (Figure 1), C_red1_ and C_red2_ are EPR active, with potential spectral overlap from all three clusters. For example, in RrC_red1_, ^32^ broad signals have been observed which are attributed to dipolar and exchange couplings between the B- and D-clusters. Here, we characterize CODH-II chemically poised at two potentials, making use of electron mediators ^31,34^ to facilitate reduction. Benzyl viologen (BV) for C_red1_ and triquat for C_red2_ were necessary to reduce the C cluster (see SI Section I). In the absence of these mediators, no C cluster signal was observed (Figure S2).

At -370 mV, 50% of BV is reduced, poising C cluster in the C_red1_ state which is demonstrably competent in CO oxidation. ^17^ At 20 mW, no signal is observed, but increasing the power to 46 mW reveals two C cluster conformations convoluted with a reduced B cluster component representing approximately 34% of the spectral intensity Figure 3a. While the reduction potential of the B cluster in CODH-II has not been established, it has been measured at –440 mV in *N. thermoacetica*, ^18^ and, while not formally measured in *R. rubrum*, it has been observed at –250 mV. ^32,35^ Given a DTT+BV reduction potential of ∼ –370 mV at pH 8, we use the Nernst equation to estimate a B cluster reduction potential of approximately –390 mV. The two C cluster conformations are best simulated with g = 1.97, 1.85 and 1.63 and the other with 2.00, 1.81 and 1.74, occurring in a 3:1 ratio. All spectral simulations are presented in Figure S15, with corresponding parameters in Table S2. A preliminary analysis of these two conformations is presented in the SI Section III.

**Figure 3.**
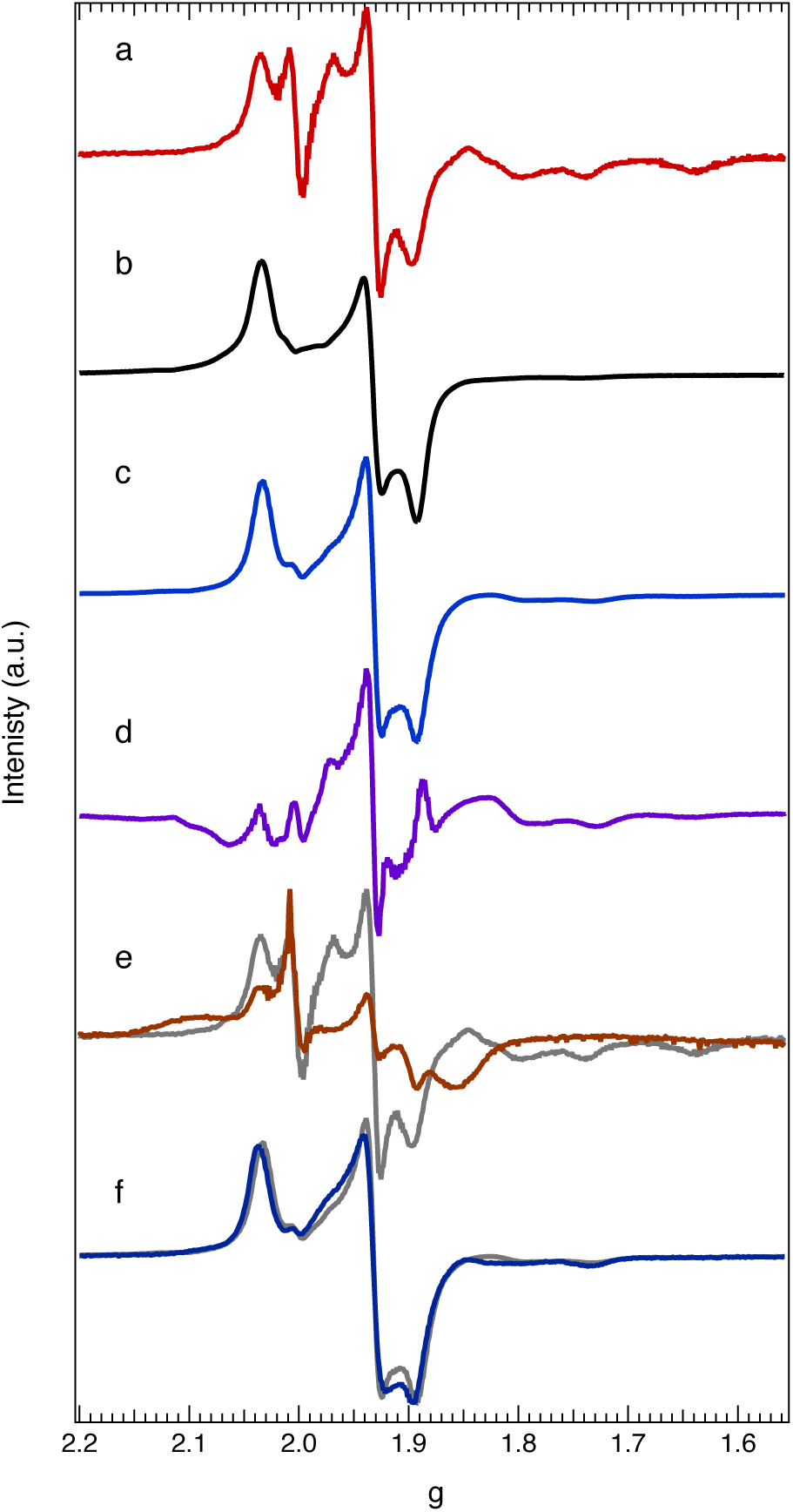
X-band EPR of CODH-II. (a) EPR spectra showing the signals acquired at -370 mV (C_red1_); (b) -430 mV (B cluster); (c) -540 mV (C_red2_); (d) The -430 mV subtracted C_red2_ spectrum highlighting the D cluster contribution; (e) C_red1_ with CN^−^ bound), C_red1_is shown for comparison in grey; (f) C_red2_ with OCN^−^ bound, C_red2_ is shown in grey for comparison. The g = 2.00 signal observed at C_red1_ is from benzyl viologen. The g values for each cluster are tabulated in **Table S2.** All spectra were collected at 10 K, other EPR parameters are available in the Methods.

At -430 mV in the absence of triquat, a rhombic signal is observed with g = 2.04, 1.93, and 1.89 (Figure 3b). This is assigned to the B cluster in its [Fe_4_S_4_]^1+^ state, consistent with Svetlitchnyi *et al.*. ^34^ Spin quantitation indicates ∼100% occupancy of the B cluster. In the absence of a mediator, a minor percentage of the C cluster is reduced, with only the g3 = 1.74 being resolvable. Simulation parameters are presented in Table S2. The D cluster (E°= ∼ – 530 mV) remains diamagnetic at -430mV. ^9,35,36^

At -540 mV, the C_red2_ state^18,34^ is observed with spectral superimposition from the B and D clusters (Figure 3c). Subtraction of the B cluster reveals the presence of two distinct rhombic signals (Figure 3d). The first, with g = 2.00, 1.95 and 1.86 is assigned to the reduced D cluster based on comparison with *R. rubrum*. ^32^ This is the first reported observation of the EPR spectrum of the D cluster for CODH-II. The similarity of the g-values between the B and D clusters in CODH-II is consistent with electronically and structurally similar [Fe_4_S_4_]^2+/1+^ sites. The remaining signal, with g = 1.97, 1.81 and 1.73, is assigned to the substrate-free C_red2_ state and is comparable to that from *N. thermoacetica*,^19^ which also displays a gz≫gy>gx spectrum consistent with a distorted T-shape geometry about the Ni center. ^37^

C_red1_ and C_red2_ were incubated with CN^−^ (CO analogue for C_red1_) and CNO^−^ (CO_2_ analogue for C_red2_), yielding significant shifts in the EPR spectra. In the presence of CN^−^, the C_red1_ signal shifts downfield to g = 2.01, 1.88 and 1.85 (Figure 3e), and a broad feature at g = 2.10 appears, consistent with dipolar and exchange couplings between the B and D clusters. ^32^ CNO^−^ binding to C_red2_ has a more modest impact, with g values of 1.97, 1.85 and 1.73 (Figure 3f). These data further substantiate the resolution of individual EPR signatures for the mediators (a sharp signal at g=2.0), B, D, and C clusters.

## 3. Fe K-edge XAS Reveals Sequential, C-Cluster-Localized Reduction

Fe K-edge XAS spectra were collected on the C_ox_, C_red1_, and C_red2_ states, representing the signal from all five iron-sulfur clusters in the CODH-II dimer (Figure 4a). The spectra show a progressive red shift with each reduction step. Fits to the pre-edge region reveal a total shift of ∼0.5 eV, while the rising edge shifts by a total of ∼0.4 eV from C_ox_ to C_red2_. The pre-edge amplitude also decreases by ∼11% with each successive reduction (Table S3, Figure S3b). Parallel Fe Kα HERFD-XAS measurements confirmed these trends and revealed a prominent split in the pre-edge region, which was used to guide the two-peak fits of the higher signal-to-noise conventional XAS data (Figure S4, Table S4).

**Figure 4.**
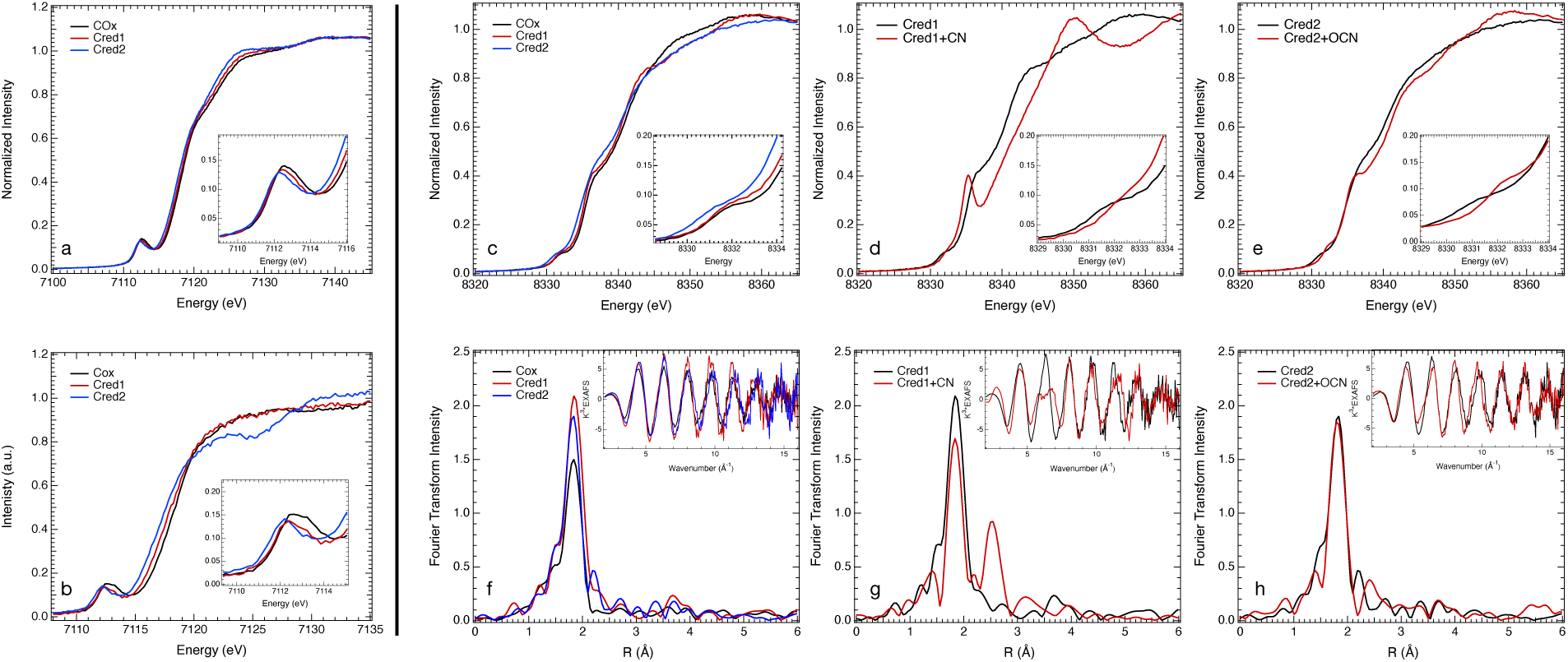
Fe K-edge XAS (a) for the three CODH intermediates under consideration. (*inset*) The expanded K-pre-edge region highlighting the 1s→3d transition. (b) EPR-informed subtractions of the CODH Fe data providing an estimate of the C cluster Fe K-edge signal. c) Ni K-edge XAS for the three CODH intermediates under consideration. Ni K-edge XAS of d) C_red1_ with and without CN^−^ bound and e) C_red2_ with and without OCN^−^ bound (*inset*) The expanded K-pre-edge region highlighting the 1s→3d transition. Ni EXAFS and corresponding Fourier transform of f) the substrate-free intermediates, g) C_red1_ with and without CN^−^ bound and h) C_red2_ with and without OCN^−^ bound.

While the B and D clusters are diamagnetic in C_ox_, approximately 34% of reduced B cluster was observed in the C_red1_ EPR spectrum. Therefore, the initial energy shift observed between the two states must arise from the one-electron reduction of Fe sites within the C-cluster, specifically the [Fe_3_NiS_4_] cubane core, as Fe_u_ is expected to remain redox-inactive, in addition to fractional reduction of the B cluster. To isolate the spectral signature of the C-cluster, we used *Clostridium pasteurianum* ferredoxin (*Cp*Fd) - a classic model for biological [Fe_4_S_4_]^2+^ clusters,^38–40^ as a spectroscopic proxy for the B and D clusters. This choice is justified by the close parallels between the two systems: the reduction potentials for the two clusters in *Cp*Fd (-347 mV and - 533 mV) ^41^ closely match those of the B and D clusters in CODH-II, and their reported EPR g-values are similar to those measured here. ^42–44^ Critically, Fe K-edge XAS data on *Cp*Fd show that its pre-edge (at ∼7112.4 eV) and rising-edge (at ∼7111.9 eV) encounter no significant shift upon [Fe_4_S_4_]^2+/1+^ reduction, while the pre-edge amplitude decreases slightly (Figure S5). This spectral invariance is attributed to a minimal structural perturbation on reduction, characteristic of clusters evolved for electron transfer, and allows for a semi-quantitative subtraction of the B and D cluster contributions from the experimental spectra. These subtractions account for oxidized B and D clusters in C_ox_, 34% reduced B cluster and fully oxidized D cluster in C_red1_ and reduced B and D clusters in C_red2_. The resulting C-cluster-only spectra and relevant parameters are presented in Figure 4b and Table S3, respectively. They show the same trends but with the energy shifts between states amplified.

The pronounced edge shift in the C-cluster data is distinct from that of typical ferredoxins, indicating significant electronic perturbation at the Fe sites upon reduction. As direct structural and functional models of the C-cluster cubane are unavailable for comparison, we turned to a series of well-characterized synthetic biomimetic cubanes, K_n_[Fe_4_S_4_(DmpS)_4_] (n=0-2), ^45–47^ which lack the structural stabilization conferred by a protein matrix and unlike the related ferredoxins, large, quantifiable Fe K-edge shifts were reported upon sequential reduction. They provide an excellent benchmark for Fe-centered reduction. The total rising-edge shift of ∼0.77 eV between C_ox_ and C_red2_ closely matches the ∼0.84 eV shift observed between the +2 and +0 states of the model series. A similar strong correlation is observed in the pre-edge: a shift of ∼0.5 eV for the C-cluster vs. ∼0.6 eV for the model compounds (Table S3). Furthermore, a nearly 40% decrease in the pre-edge amplitude is also observed for both systems, though a direct comparison of intensities is complicated by structural dissimilarities which impact the extent of 4p mixing. Thus, the trends in the Fe K-edge XAS of C_ox_, C_red1_ and C_red2_ in comparison with the models provide compelling evidence that two sequential, one-electron reductions localized on the Fe component of the C-cluster cubane occurs across this series.

C_red1_ and C_red2_ were incubated with the CO and CO_2_ mimics CN^−^, and OCN^−^, respectively and their Fe XAS are presented in Figure S6. No changes were observed in the rising edge or pre-edge upon CN^−^ binding to C_red1_, whereas OCN^−^ binding resulted in a small reduction in pre-edge intensity. These results are rationalized and consistent with the Ni K-edge XAS and EXAFS presented below.

## 4. Ni K-edge XANES Assigns a Ni^1+^ State to the Fully Reduced C-Cluster

Ni K-edge XAS data were measured for the C_ox_, C_red1_, and C_red2_ states and are presented in Figure 4c-e and Figure S7. The spectra contain two key regions that report on the Ni electronic and geometric structure. To lower energies, a weak pre-edge feature is present, which is assigned to the formally electric dipole-forbidden, quadrupole allowed Ni 1s→3d transition. The pre-edge is a sensitive probe of electronic structure, typically shifting to higher energies with an increase in ligand field strength, a pattern generally associated with an increase in oxidation state and gaining intensity through symmetry-mediated 4p mixing into vacant 3d orbitals. ^48,49^ Least-squares fits to the pre-edge show that the peak position is identical for C_ox_ and C_red1_ at ∼8331.9 eV and significantly red-shifted (∼8331.3 eV) in C_red2_ (Table 1, Figure S3). The intensity of this pre-edge feature remains nearly constant across the series.

**Table 1.** XAS parameters.

| **Table 1. XAS parameters** |  |  |  |  |
| --- | --- | --- | --- | --- |
| Nickel | Edge position at $\mu E = 0.5$ | Pseudo-Voigt Centroid | Pseudo-Voigt Amplitude | $\chi^2$ |
| Cox | 8339.17 | 8331.9 | 0.080 | 4.31E-05 |
| Cred1 | 8338.81 | 8331.9 | 0.078 | 9.42E-05 |
| Cred2 | 8337.91 | 8331.3 | 0.069 | 5.48E-05 |
| Cred1+CN | 8340.54 | 8332.7 | 0.114 | 6.22E-04 |
| – | Peak 2 | 8335.2 | 0.749 |  |
| Cred2+OCN | 8339.28 | 8332.4 | 0.116 | 9.24E-05 |
| Corresponding fits are presented in Figure S3 |  |  |  |  |

To higher energies, the rising edge position responds to changes in the coordination number, coordinating atom type, and formal oxidation state of the absorbing metal. ^50,51^ For a given series with comparable ligand systems such as substrate-free C cluster Ni, the position predominately reflects the oxidation state. Furthermore, the presence and intensity of shoulders in the rising-edge are sensitive to the local geometry, where an increase in intensity can reflect greater linearity of the ligand-metal complex, with square planar, T-shaped and linear complexes typically exhibiting sharper rising edge features. ^50,51^ The rising edge undergoes a total shift of ∼-1.3 eV from C_ox_ to C_red2_, consistent with approximately one-electron reduction. ^50,51^ Crucially, the magnitude of this shift is unequal, with the larger shift occurring between C_red1_ and C_red2_. Finally, the shoulder feature on the rising edge remains similar for C_ox_ to C_red1_, before broadening and increasing in intensity in the C_red2_ state. A useful comparison for the Ni K-edge XAS data of C_ox_ is with the A cluster in acetyl CoA synthase (ACS). In the A_ox_ state, the Ni_p_ is coordinated by three cysteine-thiolate ligands and a weakly bound water in an approximately square planar geometry, resulting in a distinct rising edge shoulder like the CODH intermediates (Figure S8). This shoulder sharpens when the water is replaced by a tightly binding methyl group, forming A_Me_. The lack of distinctive sharpening suggests the lack of a tightly bound 4^th^ ligand in the CODH intermediates, beside the three S ligands from the cubane and S(Cys526) (Figure 1). The invariance of the pre-edge energy and intensity position and the small ∼0.4 eV red shift in the rising-edge provides compelling evidence that the one-electron reduction between C_ox_ and C_red1_ is not Ni-centered and that the Ni ion remains in the Ni^2+^ oxidation state in both species. The modest rising-edge energy and intensity differences are likely due to a weak water ligand or perturbation in the Ni-S distance (see EXAFS below). In contrast, the red-shift of the pre-edge and the larger ∼0.9 eV shift in the rising-edge between C_ox_ and C_red2_ are strongly indicative of a Ni^1+^ center reduction in C_red2_.

### Cyanide binding to C_red1_

Dramatic changes are observed in the Ni K-edge when C_red1_ is treated with the CO-competitive inhibitor CN^−^ (Figure 4). The pre-edge blue-shifts by ∼0.8 eV consistent with a short Ni-CN bond and a concomitant increase in ligand-field. Furthermore, a sharp rising edge shoulder is observed at ∼8335.2 eV. While this can be partially attributed a more square planar geometry at Ni, its intensity confirms the formation of a linear Ni-CN bond with strong Ni-CN π* back bonding character as opposed to a bent binding mode with less overlap, both of which have been previously observed crystallographically. ^12,14,25^ Electrochemical data have demonstrated that CN^−^ (unlike CO) binding to the C_red1_ state does not lead to catalytic turnover. ^16^ Thus, CN^−^ binding reflects a large increase in ligand field and strong-linear coordination while the C-cluster remains in the C_red1_ redox state (see Figure 1).

### Cyanate binding to C_red2_

Incubation of C_red2_ with OCN^−^, a structural analog of CO_2_ typically used as an inhibitor, ^13^ allows the trapping of a ligand-bound C_red2_ state which has been investigated previously using electrochemical methods. ^16^ Ni K-edge XAS data show dramatic changes upon OCN^-^ binding to C_red2_. The pre-edge blue-shifts by ∼1.0 eV consistent with a short Ni-C(ON) bond and a concomitant increase in ligand-field. The rising-edge shows a shoulder consistent with square-planar geometry. While evidence of slow turnover of the OCN^−^ to the CN^−^ bound form has been reported, ^52^ The lack of an intense rising-edge feature suggests a non-linear Ni-C(N)-O geometry and little to no turnover of the OCN^−^ to CN^−^ during the 15 minute incubation time. Thus, these data suggest a bent OCN^−^ coordination to the Ni center and the formation of a strong Ni-C(ON) bond.

## 5. Solution EXAFS Clarifies Crystallographic Ambiguity in C-Cluster Coordination

Despite numerous crystallographic studies on CODH-II, the precise structure of the C-cluster remains contentious, partly due to the enzyme’s extreme oxygen sensitivity which can lead to cluster decomposition. ^15^ Six substrate-free CODH-II crystal structures have been solved (PDBs 1SUF – inactive protein; 1SU7, 3B53 and 9FPG-Cred1, 1SU8 and 3B51 – Cred2) (see SI Section IV for a detailed comparison). The quality of all structures was assessed via calculated RSZD scores ^53^ as implemented in CCP4 ^54^ (Table S5). The RSZD score is a measure of crystal model accuracy; values exceeding a cutoff of 3σ represent significant atomic misplacement (for negative σ values) or unexplained electron density (for positive σ values). The RSZD scores for intact C clusters across available CODH crystal structures were assessed revealing a median value of 6.6σ. This highlights the difficulty in structurally modelling the C-cluster. The analysis (see SI Section V) reveals that in all six cases, the atoms of the [Fe_3_NiS_4_-Fe_u_] cluster exceed this threshold. While 1SUF, 1SU7, and 1SU8 diverged the most, 3B53 and 3B51 exhibited a) substantial unexplained density at the Fe_u_ site, b) misplaced atoms and c) unexplained density within the [Fe_3_NiS_4_-Fe_u_] pseudo-cubane. Notably, even the recent high-resolution C_red1_ structure (9FPG) exhibits substantial atomic misplacement in the cluster (-7.9σ) and critically, at the key Ni-coordinating residue Cys526 (-7.1σ). Figure S9 highlights the overfitting in the Cys-Ni bond in 9FPG and indicates that the Ni is in two alternative configurations which are not explained. This is also observed in the accompanying CO-CO_2_-water bound structure 9FPO, underscoring the uncertainty in the Ni position, and inaccuracy of the Ni-L and Ni-Fe bond lengths as these. Bond lengths from each structure are presented in Table S6 for comparison to the EXAFS presented below, and an analysis of the fitting of these Ni configurations in these structures is presented in SI Section IV. This indicates that while the available crystal structures provide a valuable overall scaffold, they are compromised in their accuracy for defining the precise ligand identities and metal-ligand distances at the C-cluster active site, despite the high resolution of the overall structure. These issues suggest the need for additional methods, as described here, to assign the structures and electronic states of the intermediates in the CODH mechanism.

While the precise metrics differ, crystallographic studies consistently show the Ni center integrated into the cluster via two sulfide ligands and one cysteine ligand (Cys526) from the protein, forming an open cubane, as shown in Figure 1. This has led to a consensus model of a persistent trigonal Ni coordination. ^10,11,14,25^ Starting from this understanding, our solution-state Ni EXAFS data reveal important structural differences between the three redox states that are not resolved by crystallography. The k^3^-weighted Ni EXAFS were measured to 16 Å^-1^ and their corresponding Fourier transforms (FTs) were computed for the three C-cluster intermediates (Figure 4f). The data show significant changes in phase and amplitude upon successive reduction.

FEFF-based first-shell least-squares fits (Table 2, Figure S10a,b, Table S7) reveal that, while C_ox_ can be modelled with 3 single Ni-S distances at 2.22 Å, a split first shell with 2 Ni-S 2.20 and 1 at 2.30 Å provides significant improvement (Table S7). We therefore propose a distorted three coordinate site for C_ox_. Upon one-electron reduction to C_red1_, the FT first shell intensity increases, and the Ni site remains three-coordinate with the 3 Ni-S distances best fit at 2.22 Å. This indicates that upon reduction of the cubane portion of the C-cluster, the Ni-S distances become more ordered, and the average Ni-S becomes shorter. In the C_red2_ state, a further modest contraction of the 3 Ni-S to 2.19 Å is observed. Note that, despite this shortening, the Ni K-pre-edge for C_red2_ exhibits a ∼0.6 eV red-shift relative to C_red1_, providing further support on a Ni-based reduction from C_red1_ to C_red2_. Note that while the first shell is dominated by S ligands, weaker, longer water coordination cannot completely be ruled out (see Table 2, Table S7 and Figure S10). Furthermore, there is little intensity in C_ox_ beyond the first shell, which increases in C_red1_ and C_red2_. The structural implications are discussed below.

**Table 2.**
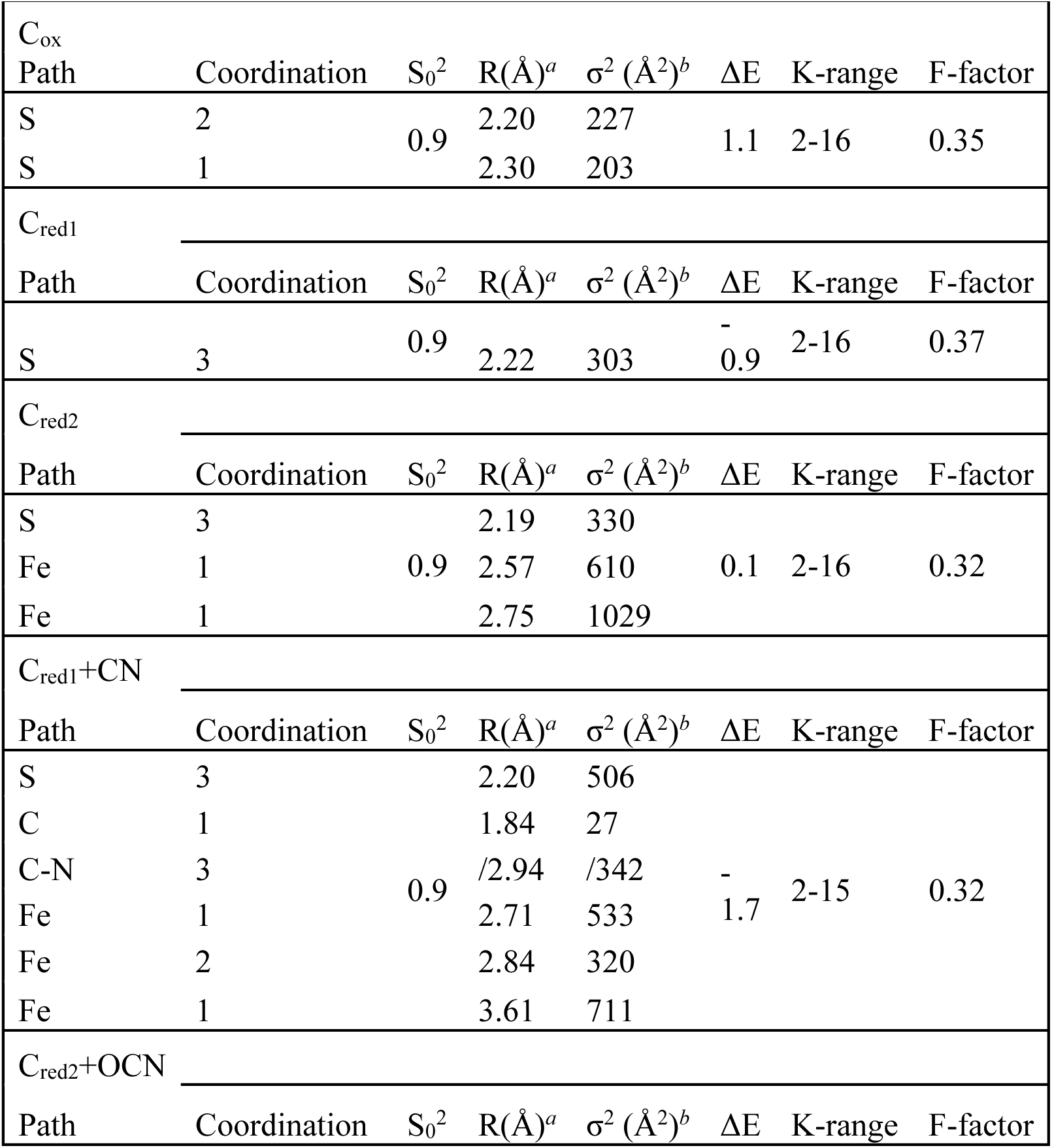
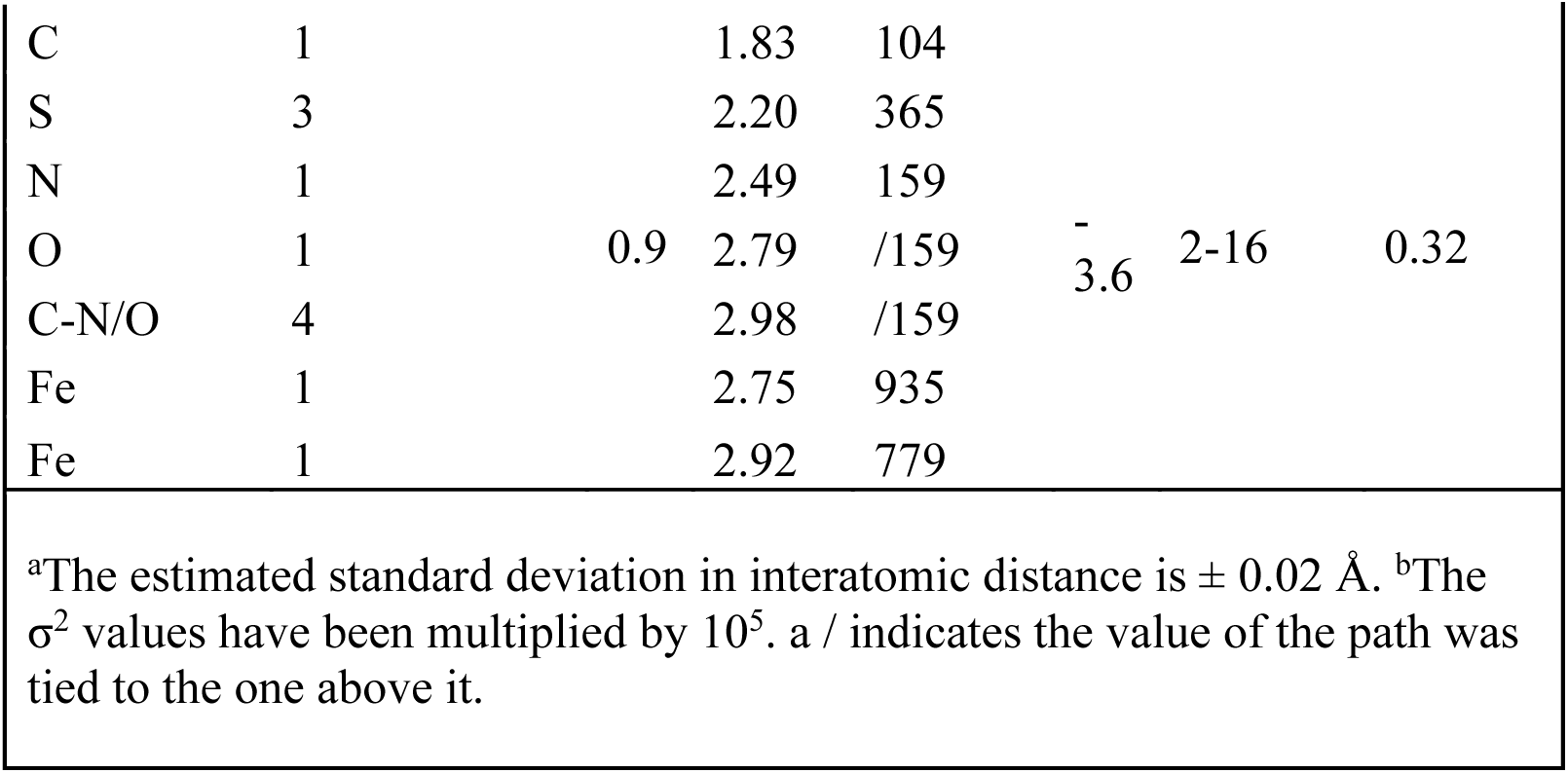
EXAFS least-squares fitting parameters.

## 6. Substrate Analogues Bind to C_red1_ and C_red2_ and Establish Dynamism in Cluster Ordering

Incubation with substrate analogues CN^−^ and OCN^−^ profoundly impacts the Ni-EXAFS of C_red1_ and C_red2_ (Figure 4g,h), respectively. These data along with the EPR and Ni K-edge XAS on the substrate bound species, support the clean conversion of C_ox_ to C_red1_ to C_red2_. First shell least-squares fitting of the C_red1_+CN^−^ EXAFS shows a Ni-C at 1.84 Å and 3 Ni-S at 2.20 Å. Fitting of the C_red2_+OCN^−^ data reveal a short Ni-C at 1.83 Å and 3 Ni-S at 2.20 Å. The corresponding multiple-scattering paths from a linear Ni-CN (in C_red1_+CN^−^) and a bent cyanate (for C_red2_+OCN^−^were required to model the data. Interestingly, despite inclusion of these components, the FT show unfit intensity in the second shell that are consistent with prominent Ni-metal interactions. A final fit for C_red1_+CN^−^ includes Ni-Fe components at 2.71 Å, 2.84 Å and 3.61 Å (Table 1, Table S7, Figure S10). Similarly, the C_red2_+OCN^−^ data show Ni-Fe components at 2.75 Å and 2.92 Å.

The observation of Ni-metal components in the EXAFS of the substrate analogue-bound species allows us to explain a persistent mystery in the EXAFS of CODH in the literature: the low intensity, or absence of features in the second shell of the FT, where more intense Ni-metal components should be observed. ^28–30^ This has been attributed to structural disorder and associated dampening of these signals. ^29^ This explanation is consistent with data on 4Fe-4S clusters where increased disorder in Fe-Fe distances leads to significant damping of the Fe-Fe FT feature with reduction via increasing the EXAFS Debye-Waller term. ^45,55^ The lack of Ni-metal FT components has also been attributed to destructive interference between multiple Ni-Fe components. ^29^ FEFF simulations (Figure S11) indicate that two Ni-Fe paths with similar Debye-Waller terms (a measure of static and thermal disorder) are maximally out of phase at differences of ∼ 0.2 Å, destructively interfering to modulate their contribution to the EXAFS signal close to the noise level. Small deviations from this difference are enough to modulate the degree of destructive interference significantly (Figure S11). This relationship is discussed further in SI Section VI. The two closest Fe atoms to the Ni center, Fe2 and Fe_u_ (from consensus crystallography, Figure S1, Figure 2, Table S1), are expected to contribute to the EXAFS. Interestingly, the mean crystallographic Ni-Fe distances are 2.70 ± 0.25 (Fe_u_) and 2.86 ± 0.22 (Fe2) – a separation of ∼0.2 Å (Figure 2). We propose that the prominence of Ni-metal contribution in the substrate bound forms (Figure 4g,h) and in C_red2_ (Figure 4f) occurs when their difference deviates from ∼0.2 Å given equal Debye-Waller terms. Comparatively, CN^−^ binding to Ni yields a small but meaningful change in the Ni-metal distances relative to C_red1_ (ΔFe-Fe = 0.04 Tables S7, S6-Cred1-g, S8), while having much lower Debye-Waller terms, indicating a lower disorder and larger EXAFS contribution.

Using the average crystal structure distances as starting models, second shell fits were performed on all 5 species (Table S7, Figure S10c-i). The fits show that the Ni-Fe components become prominent as their difference deviates from ∼0.2 Å, resulting in a stronger cancelling effect for C_ox_ and C_red1_. Note that disorder also plays a role in the damping since the Debye-Waller terms are also different across these fit components. While the Ni-metal contributions in C_ox_ and C_red1_ are weak, the fits to the other three species were used to inform and constrain the in initial distances in the fitting procedure, increasing the confidence in the Ni-Fe components in these species.

The available PDB data (Figure 2) shows that the Ni-Fe_u_ distance is consistently shorter than the Ni-Fe2 distance, providing a basis for the attribution of the shorter and longer Fe measured as Fe_u_ and Fe2, respectively. Fe_u_ in C_ox_ and C_red1_ is at comparable distances, whereas Fe2 shows more variability, contracting upon reduction of C_ox_, suggesting compression of the rhomb formed by Ni-S_2_-Fe2. This is consistent with compression of the Fe_2_S_2_ rhombs observed in typical [Fe_4_S_4_] clusters. ^45,47^

Further reduction to C_red2_ gives a Ni-Fe_u_ distance of 2.57 Å, which positions Ni and Fe_u_ for CO_2_/OCN^−^ coordination. Ni-Fe2 shortens by ∼0.13 Å, again consistent with compression of the Ni-Fe2-S_2_ rhomb as the C cluster gains two additional electrons. Cyanate binding elicits a more substantial change, with Fe_u_ and Fe2 elongating by approximately the same amount (∼0.18 Å). Elongation of Fe_u_ after OCN^−^ binding indicates that the close Ni-Fe_u_ distance is only required to initially bind the substrate, with Fe_u_ likely coordinating the O(CN)^−^, made possible by a distortion of OCN^−^ from linearity. ^13^ The elongation of Fe2 suggests perturbation of the Ni-Fe2-S_2_ rhomb likely the result of structural rearrangement to accommodate the Fe_u_-O(CN) bond and the simultaneous formation of a tight Ni-C(ON) bond (Ni-C ∼1.83 Å).

## 7. An Integrated Spectroscopic Model

The combined spectroscopic data allow for a definitive assignment of the electron distribution within the C-cluster across its catalytic substrate free states. The red-shifts in the rising edge for the 3-electron reduction from C_ox_ to C_red2_ (∼1.3 eV in the Ni K-edge and ∼0.8 eV in the Fe K-edge) are consistent with a highly delocalized cluster where sequential reduction impacts the entire [Fe_3_NiS_4_-Fe_u_] complex. The data presented here support the formal assignments of the C-cluster states as follows: C_ox_: Ni^2+^[Fe_3_S_4_]^2+^-Fe_u_^2+^, C_red1_: Ni^2+^[Fe_3_S_4_]^1+^-Fe_u_^2+^, C_red2_: Ni^1+^[Fe_3_S_4_]^0^-Fe_u_^2+^. This is consistent with the EPR assignments: diamagnetic C_ox_ and paramagnetic C_red1_ and C_red2_.

The assignment of the iron-sulfur component in the C_red2_ state as an all-ferrous [Fe_4_S_4_]^0^ cluster is also significant, as this state is rare in nature. The most notable biological example is the nitrogenase iron protein (FeP). ^56^ Given the rhombic EPR spectra obtained for the S=1/2 C_red2_ state, it is plausible that the integer-spin all-ferrous [Fe_4_S_4_]^0^ component is in an S=0 ground state, analogous to FeP, which can adopt either a S=0 or S=4 catalytically competent ground state ^57^. Our XAS data also suggest a high degree of delocalization within the formal assignment of Ni^1+^-[Fe_4_S_4_]^0^ (C_red2_) and Ni^2+^- [Fe_4_S_4_]^1+^ (C_red1_) with the edge shifts suggesting spin delocalization between the Ni and the [Fe_u_-Fe_3_S_4_] component in both reduced species. This may also be responsible for the similarity in the g values obtained for the two species. Careful computational investigation of the multiconfigurational ground state of the C-cluster, in addition to DFT-based simulations of the EPR and XAS data are necessary to better understand the role of delocalization and spin coupling and will be the focus of a future study. We note that the current extent of computational work on the C cluster is limited to a handful of studies primarily investigating energetics and mechanism development, and lacking correlative spectroscopic data. ^23,58,59^

## 8. Contrasting with Alternative Models for the C_red2_ State

It has been proposed that both electrons in the reduction from Cred1 to Cred2 are localized on the nickel, yielding a Ni^0^ state. ^21^ However, as noted previously, the ligand environment isn’t appropriate to stabilize this low valent species. ^22^ More definitively, our Ni K-edge XAS data are inconsistent with a d^10^ Ni^0^ center, both in the magnitude of the rising-edge shift and in the existence of a clear 1s→3d pre-edge transition. This finding mirrors our previous results for the A-cluster, where too a Ni^0^ state has been unobserved, with substrate (e.g. CO) binding promoting reduction to a Ni^1+^ state. ^60^ Another proposal rooted in computational studies on the crystal structures from Jeoung and Dobbek 2007 ^11^ involves the formation of a Ni^2+^-hydride species. ^23^ This has been challenged by later computational work demonstrating such a pathway would likely yield a protonated cubane sulfide, and eventual loss of the hydride via H_2_ formation ^59^ or generation of formate instead of CO by a hydride transfer. ^61^ Experimentally, we have demonstrated that CODH-II (like the N. thermoacetica CODH) has very low H_2_ evolution activity, whereas Ni-hydride species are expected to evolve adventitious H_2_. ^62–64^ The lack of hydrogenase activity (H^+^/H_2_, E^0’^ = -420 mV) is one of the most interesting features of CODH (CO_2_/CO, E^0’^ = -540 mV) and is an important attribute of a CO_2_ reduction catalyst. Most CO_2_ reduction catalysts suffer from competing proton reduction, which significantly lowers their efficiency. Further, metal-hydride bonds are known to cause significant perturbations to the metal XAS rising-edge region while having minimal diagnostic impact on the EXAFS. ^51,65^ Such behavior is not observed between our C_red1_ and C_red2_ spectra. Another proposal by Lindahl et al. argues that the electrons are stored in a dative Ni-Fe bond, which is postulated to have minimal impact on the spin system, providing a tidy explanation for the similarity observed in the EPR signals for C_red1_ and C_red2_ in other CODH orthologs. ^22^ This has gained support from studies by Holland et al. wherein C-cluster structural analogues containing a W atom were synthesized. ^66^ Upon reduction, electrons were found to be stored in a M–M σ bond between Ni and the cuboidal W. However, the relatively diffuse 5d manifold of W may facilitate σ-bond formation in ways that the contracted 3d manifold of Fe is unable to replicate in the C-cluster. In a follow-up study by the same investigators, clusters containing only Fe and Ni were synthesized. ^67^ They found that the addition of Ni as Ni^0^ resulted in a Ni^1+^ and reduced [Fe_4_-S_4_]^0^ cluster, wherein the added electrons were preferentially stored on the Fe sites, as opposed to within a dative Ni-Fe bond. Our XAS results on the C-cluster are consistent with these latter findings. Finally, a recent investigation by Basak et al. observed that the two-electron reduced C_int_ state accumulated at the same negative potential as C_red2_ in the absence of substrate, with subsequent conversion to C_red2_ state after exposure to CO_2_. ^20^ This led to a proposal that the reducing equivalents required to form C_red2_ are stored in a carbonite-type Ni-CO_2_ interaction. Our data directly contradict this model. We observe (via EPR) the quantitative formation of the substrate-free C_red2_ state in the absence of CO_2_, and our Ni and Fe K-edge XAS data demonstrate that the reducing equivalents are stored within the metal cluster, forming the Ni^1+^-[Fe4S4]^0^ state.

## 9. Resolving Discrepancies in the CODH-II Literature

Our findings clarify several conflicting reports in a recent study by Basak et al. on CODH-II. ^20^ A key point of divergence is the nature of the C_red1_ state. Basak et al. present an EPR signal for a Ti(III)-EDTA reduced C_red1_ (g = 1.97, 1.82, 1.64) obtained via subtraction from the substantial Ti(III) background signal. However, the data provided with the study suggests detector saturation from the Ti(III) signal, making quantitative subtraction infeasible (Figure S12). Furthermore, this sample was poised at -409 mV, comparable to the -430 mV potential at which we observe quantitative B-cluster reduction. Indeed, the spectrum reported by Basak et al. appears to contain features consistent with the reduced B-cluster signal reported here.

A second major discrepancy lies in the formation of the C_red2_ state. Using strong reductants alone, Basak et al. were unable to observe quantitative C_red2_ formation in the absence of CO_2_.^20^ Additionally, they assert that CO_2_ binding is the key distinguishing feature that defines the C_red2_ state. This conflicts with our results, which show a robust, substrate-free C_red2_ EPR signal. It also is at odds with extensive electrochemical and EPR studies of CODH, which demonstrate that CO_2_ binds to the C_red2_ state. ^16,19^ The critical difference in our experiments was the use of an organic redox mediator (triquat) to facilitate electron transfer from the reductant to the D cluster. We observed poor yields of both C_red1_ and C_red2_ in the absence of mediators (as diagnosed with Ni K-edge shifts and EPR) (Figures S2, S13). Our results suggest that the use of a suitable mediator is required for conversion to the substrate-free reduced states of the C-cluster, agreeing with prior work in CODH-II ^34^ and recent work in RrCODH. ^31^ Finally, our resolution of the B-and D-cluster EPR signals helps to reinterpret previously published spectra. At -530 mV, Basak et al. report g-values for the reduced B-cluster (g = 2.03, 1.93, 1.90) ^20^ which appear to be a mixture of the g-values which we have now disambiguated for the separate B- and D-clusters.

This suggests the spectrum obtained at this potential is a composite of undistinguished signals from both clusters.

## 10. The Mechanism

The consensus mechanism of CODH as understood prior to the recent work by Basak et al., ^20^ is summarized in Figure 1, and incorporates the results of this study. Basak et al. ^20^ modify this mechanism by stipulating that: C_red1_ contains a hydroxyl bridging Fe_u_ and Ni, CO actually binds to a C_int,o_ species and not to C_red1_, in C_int_ Fe_u_ is bound to a hydroxyl, and C_int,i_ is the CO_2_ binding species, not C_red2_. Our data does not support the mechanistic development proposed by Basak et al., favoring instead the consensus model, while expanding it with the oxidation state assignments and structural changes reported here. Notably, in C_red1_, we see no evidence of a hydroxyl bound to Ni in a bridging fashion or otherwise, and we observed CN^−^ binding to and stabilizing the C_red1_, with no evidence of reduction to C_int_. While C_red2_ does contain a short Ni-Fe_u_ distance (2.56Å), which is required for substrate binding, we see no evidence of the short Ni-Fe_u,i_ distances proposed by Basak et al., which range from the improbable 2.07 Å to the more likely 2.60 Å in their study (Table S9). Perhaps most importantly, we demonstrate that C_red2_ is the species competent to bind the CO_2_ substrate analogue OCN^−^ and does not require the addition of substrate to form, as proposed by Basak et al. ^20^

Beyond the consensus model, our data establish key oxidation states and bond lengths for the catalytic intermediates of CODH. We observe similar Ni-Fe_u_ distances in C_ox_ and C_red1_, which contract by 0.1 Å on reduction to C_red2_, demonstrating that the two-electron reduction elicits a structural rearrangement of the cluster in preparation to bind CO_2_, bringing the Fe_u_ and Ni into close quarters to adequately coordinate the C and O. On binding of CO_2_, the Ni and Fe_u_ expand, allowing for Fe_u_’s role in stabilizing O for protonation, a critical step in the reduction of CO_2_. Similarly, CN^−^ binding to the C_red1_ state elicits a substantial rearrangement of the cluster, as it responds to ligand binding by becoming more ordered in a way detectable by EXAFS, that is the distribution of Ni-Fe distances becomes such that the competing Ni-Fe paths no longer destructively interfere to a substantial degree. These observations are all consistent with the consensus mechanism. ^14,58,68^

## 11. Conclusion and Summary

Here, we have used a robust protocol for the expression, purification, and high-yield trapping of CODH-II from *C. hydrogenoformans* in its C_ox_, C_red1_, and C_red2_ states to probe the geometric structure around the Ni center and electronic structure of the [NiFe_3_S_4_]-Fe_u_^2+^, C-cluster. The use of a redox mediator is critical for achieving complete conversion between these states, a methodological insight which allows us to address conflicting reports in the literature, such as the suggestion by Basak et al. that the substrate CO_2_ is required to reach the C_red2_ state. ^20^ Our analysis provides definitive redox assignments for the key intermediates, and our solution-state Ni EXAFS data complements the existing crystallography by providing a precise model for the dynamic coordination environment of the Ni site and the associated cluster rearrangements. The redox state assignments described here are significant because they place the two electrons generated from two-electron oxidation of CO on the Ni and Fe components of the C-cluster, instead of within a metal-metal bond, ^67^ in formation of a Ni^0^, ^21^ or in a coordinated hydride. ^23,59^ These results also suggest that an intricate tuning of the redox potential of the components of a heterometalic cluster offers a strategy to circumvent the need for the radical cation during CO_2_ reduction and offers increased efficiency relative to industrial catalysts. Another key finding of this work is the direct observation of an all-ferrous iron component in the C_red2_ state, a unique phenomenon among iron-sulfur enzymes, made more so by the presence of the catalytic Ni. Our description of the key C_red2_ state underscores the importance of a three coordinate Ni, devoid of water coordination, permitting facile binding of CO_2_ to the electron-rich Ni-Fe pseudocubane.

## Supporting Information

The Supporting Information is available free of charge: detailed methods, A detailed account of CODH, Preliminary analysis of the Cred1 EPR at High Power, A Brief history of substrate-free CODH-II Crystallography, Comprehensive RSZD Survey of All Available CODH Structures, Constructive and destructive Interference of Ni-Fe paths in EXAFS of the C cluster, Figures S1-S15, Tables S1-S12, additional references.

## AUTHOR INFORMATION

## Author Contributions

The manuscript was written through contributions of all authors. All authors have given approval to the final version of the manuscript.

## Acknowledgments

We thank Dr. Victor Mougel for making the XAS data on the biomimetic K*n*[Fe_4_S_4_(DmpS)_4_] series from Grunwald et al. 2024 ^45^ available for public use. The dataset can be accessed via its DOI at https://doi.org/10.3929/ethz-b-000742321. We also thank Dr. Graham George for sharing the CpFd XAS data set. The Stanford Synchrotron Radiation Lightsource is supported by the U.S. Department of Energy Office of Basic Energy Science under Contract No. DE- AC02-76SF00515. The SSRL Structural Molecular Biology Program is supported by the US Department of Energy Office of Biological and Environmental Research, and by the NIH National Institute of General Medical Sciences (including P41GM103393).

