## Supplementary Information for "Use of Substrate Analogues and X-ray spectroscopy Reveals an all Ferrous C-Cluster in CO Dehydrogenase"

Macon Abernathy,[a,1] Kareem Aboulhosn,[b,1] Christopher Ohmer,[a] Stephen Ragsdale,[b] Ritimukta Sarangi[a]
[a]Stanford Synchrotron Radiation Lightsource, SLAC National Accelerator Laboratory; Menlo Park, CA 94025, USA.
[b]Department of Biological Chemistry, University of Michigan; Ann Arbor, MI 48109, USA.

[1]M.A. and K.A. contributed equally to this work.

Corresponding Author: Ritimukta Sarangi

**This PDF file includes:**

### Section I: Methods

Expression and Purification

Several organisms are the focus of research within the WL pathway,(1) among these, CODH-II from *Carboxydothermus hydrogenoformans* (Ch) Z-2901 stands out for its high activity, industrial applications, and ability to be recombinantly expressed in *E. coli* (2). CODH-II was expressed and purified from *C. hydrogenoformans* Z-2901using previously published protocols (2). After purification of recombinantly expressed CODH-II, the protein is incubated with 1 mM rezarusin (5-fold excess relative to CODH-II) for 5 minutes and subsequently buffer exchanged into a storage buffer containing 50 mM Tris, 20% glycerol pH 8. All states of CODH-II are prepared in the storage buffer. The CODH-$C_{ox}$ state is prepared by incubating 2 mM CODH-II with 10 mM resazurin ($E^0$ = 110 mV) for 15 minutes and then freezing in LN2. The CODH-$C_{red1}$ state is prepared by incubating 2 mM CODH-II with 20 mM dithiothreitol and 1 mM benzyl viologen ($E^0$ = 370 mV) for 15 minutes, then freezing in LN2. UV-Vis measurement confirmed that half of the benzyl viologen was poised in the reduced state under these conditions. The CODH-$C_{red2}$ state is prepared by incubating 2 mM CODH-II with 20 mM dithionite and 1 mM triquat for 15 minutes then freezing in LN2 ($E^0$ = -540 mV). Protein concentration was determined using Rose-Bengal assay. Specific activity of CODH-II varied between 500-775 U/mg.

After purification of recombinantly expressed CODH-II, the protein is buffer exchanged into a storage buffer containing 50 mM Tris, 20% glycerol pH 8. The CODH-$C_{red1}$ EPR sample was prepared in the storage buffer by incubating 0.25 mM CODH-II with 20 mM dithiothreitol and 1 mM benzyl viologen for 15 minutes in the anerobic chamber before freezing with LN2 inside the chamber. The CODH-$C_{red2}$ state is prepared similarly but was incubated with 0.25 mM CODH-II with 20 mM dithionite and 1 mM triquat for 15 minutes in the chamber before freezing in LN2 within the chamber. The –430 mV sample was prepared with 20 mM dithionite in the absence of triquat and incubated for 15 minutes before freezing in LN2 in the chamber.

The CODH-$C_{red1}$-CN XAS sample is prepared by incubating 2 mM CODH-II with 20 mM dithiothreitol, 1 mM benzyl viologen, and 50 mM KCN for 15 minutes then freezing in LN2 in the anaerobic chamber. The CODH-$C_{red2}$-OCN XAS sample is prepared by incubating 2 mM CODH-II with 20 mM dithionite, 1 mM triquat, and 250 mM KOCN for 15 minutes and then freezing in LN2.

Feedback from Ni K-edge XAS was integral to the optimization of the protocol to produce C cluster intermediates which are as pure as possible, and free of contaminants (i.e. Cu) which would prevent the collection of high-k EXAFS data. The preparation protocol which was developed as a result of this optimization has produced intermediates which yield repeatable XAS spectra form batch to batch. We note that XAS screening was critical to the development of this protocol. For example, validating that $C_{ox}$ is truly oxidized is difficult, due to the diamagnetic nature of the intermediate, and the EPR-silent nature of $C_{red1}$ from CODH-II. Likewise, ensuring that $C_{red1}$ and $C_{red2}$ were formed, in absence of a substrate requires a spectroscopic probe like Ni and Fe XAS which is capable of differentiating the species on the basis of edge shifts and rising edge morphology corresponding to element-specific placement of the electrons, and any resulting coordination changes.

X-ray Absorption Spectroscopy

Prior to measurement, CODH-II samples (1-2 mM in concentration) were syringed under anoxic conditions into Delrin cuvettes sealed with 25 μm thick polyimide tape. These were flash frozen in liquid nitrogen and shipped to Stanford Synchrotron Radiation Lightsource for measurement. Ni K-edge XAS were collected on beamline 7-3, a 20-pole, 2T wiggler side-station. Energy selection was provided by a Si(220) double crystal monochromator oriented to ϕ = 90. The contribution of higher-order harmonics was reduced by detuning the monochromator by 40%, and unwanted scatter was reduced with soller slits and a 6-absorption length Co filter. Spectra were collected in fluorescence mode using a Canberra 32-element germanium solid state detector with Xspress3X electronics. During data collection, the sample was maintained at 6 K using an Oxford Instruments CF1208 LHe cryostat and closed-loop cryocooler. An in-line Ni foil was used as a reference, and the inflection point of the rising edge was calibrated to 8331.6 eV. Each spectrum was individually calibrated using its accompanying in-line Ni foil spectrum. Iron K-edge XAS were also collected on beamline 7-3 under the same conditions, but with the use of a 6-absorption length Mn filter, and with Fe calibrated to 7111.3 eV.

All channels were inspected for contamination by elastic scatter prior to averaging using a custom python script. Spectra were calibrated and normalized in Athena, and the EXAFS signal was extracted using Pyspline,(3) and fit using EXAFSPAK.(4) $E_0$ was set to 8340 eV and fitting a 5-region spline to the data with polynomial orders 2, 3, 3, 3 and 3. Feff7 (5) was used to generate theoretical EXAFS phase and amplitude parameters of scattering paths within a 5 Å radius of the Ni absorber within the starting structure (PDB: 4UDX or 5FLE) (6, 7). Four parameters were evaluated for each backscattering path in course of fitting: the interatomic distance R, the interatomic disorder $\sigma^2$, the shift in the assigned value of $E_0$, and the amplitude reduction factor $S_0^2$. Of these four parameters, R and $\sigma^2$ were allowed to float for each interaction, while the shift in $E_0$ was allowed to float but fixed to a common value among all paths of a given fit. $S_0^2$ was fixed at 0.9. Peak deconvolution was done in Larch v9.45 (8) using pseudo-Voigt functions to model the Ni and Fe K-pre-edge features.

HERFD measurements

High energy-resolution florescence detected X-ray emission spectra were collected at beamline 15-2, a state of the art undulator beamline equipped with a 7 spherically bent crystal Johann-type XES spectrometer (9). Measurements were collected at the Fe K$\alpha$ line. Energy selection was provided by a liquid nitrogen cooled double crystal monochromator equipped with a Si(311) for Fe at $\phi$ = 0. The resolving power of the Si(311) is approximately 10x greater than that of the Si(111). The beam was focused using a KB mirror system. Ge (440) crystals were used in the collection of Fe K$\alpha$ emission signal in a 1m Rowland geometry. A single element silicon drift detector was used to measure the signal. To limit signal attenuation, the emitted beam path was enclosed in a He-filled bag. The sample was maintained in a cryostat at 10 K to limit photodamage. All spectra were inspected for evidence of photo damage, which was monitored during data collection, and limited by attenuating the incoming photon flux using Al filters. Individual spectra were compared for evidence of beam damage before averaging. HERFD spectra were processed using PyMCA 5.9.2,(10) and normalized using Athena.

EPR Methods

X-band EPR measurements were collected with a Bruker ECS106 spectrometer equipped with a TE102 cavity, B – E25 magnet and cold edge CH-210N LHe cryostat. Spectra were collected under non-saturating, slow passage conditions at 10 K. Spectrometer settings are summarized in Table S10. All EPR simulations were done using Easyspin 6.0.11 (11) and Matlab 2025b. Parameters used in the simulation are presented in Table S2.

PDB Survey

The PDB was surveyed for all CODH structures. Structures were identified via a search for the C cluster ligand, which is identified using the following codes: WCC, NFS, XCC, BF8, RQM, and 82N. The resulting survey yielded 92 structures. The C clusters of each structure were extracted, and any multiple conformations were parsed into their own xyz structures on the basis of altloc identifiers in the PDB file. For example, 8CMW contains 3 conformations of the C cluster, and each was assigned to its own structure file. Two conformations of the C cluster from 6B6W were excluded, on the basis of an apparent bonding interaction between K563 (CODH-II numbering) and Ni. The resulting 186 C cluster structures were used to extract the Ni-S distances coming from Cys526, S1 and S2 (as defined in Figure S1), along the Ni-$Fe_u$ and -Fe2 distances, and the resulting data set was curated to remove outliers. Outliers were identified as any structures containing a Ni-S bond (with S coming from Cys526, S1 or S2) greater than 3 Å (26 structures), or a Ni-Fe distance shorter than 2 Å (2 structures). The remaining 158 C cluster structures were used to make Figure 2 and Table S1. Summary values are presented in Table S11.

CODH sample preparation optimization

One measure to assess the completeness of C-cluster assembly is the specific activity of the CODH. Over the course of protocol optimization, (Figure S13) specific activity was optimized according to work that was done in parallel with the work described here (2). For example, the protein used for the January 2024 measurement (Figure S13) had a specific activity of only 512 U/mg whereas the December 2024 measurement had protein with a specific activity of 775 U/mg. This increased activity owed to better controls for oxygen and reagent purity. One important control for oxygen is rapid freezing of the samples inside the anaerobic chamber rather than removing them in a secondary container for subsequent freezing in LN2 under atmosphere. Even short exposure to oxygen will rapidly degrade the C-cluster and alter the desired state of the protein (12).

Another iterative process was generating $C_{ox}$, $C_{red1}$, and $C_{red2}$; the major challenge being getting complete reduction of the clusters relative to $C_{ox}$. In earlier measurements such as October 2023 and January 2024 (Figure S13), only a 5-fold excess of reductant relative to CODH was being used. Additionally, because the CODH was purified with a reductant, the difference in oxidation states of $C_{red1}$/$C_{red2}$ compared to $C_{ox}$ were not significant. Starting in March 2024, a 10-fold excess of reductant was used to drive the protein to more complete reduction. Although this increase seemed to help distinguish the states, it was still not leading to the expected changes in the rising edge (Figure S13).

Using reductant alone does not reliably generate the $C_{red2}$ state even when used in great stoichiometric excess (Figure S2). The EPR data indicated that $C_{red2}$ was reliably generated (Figure 3c, S2) only with the addition of triquat as a redox mediator. We decided to begin incorporating mediators in XAS sample preparations in June 2024. Increasing the incubation time in July 2024 to 15 minutes with CODH, mediator, and reductant further improved reduction. However, the redox states of CODH were still not fully distinguished. We suspected that $C_{ox}$ may not be fully oxidized because the protein is always purified with reductant to protect against oxygen damage. To address this issue in December 2024, the CODH was oxidized with resazurin after purification but before buffer exchange; then following buffer exchange was treated again with resazurin. This method resulted in the fully oxidized $C_{ox}$ state on the basis of the XAS measurements.

This iterative process enables the study of $C_{ox}$, $C_{red1}$, and $C_{red2}$ by XAS. Additionally, because cyanide and cyanate selectively bind only to $C_{red1}$ and $C_{red2}$ (respectively), it also enabled the study of the electronics of these competitive inhibitors.

### Section II: A Detailed Account of CODH

At the surface of the homodimer is situated the D cluster, shared by the two monomeric units of the dimer and is solvent exposed. It is involved in delivering or expelling electrons from the clusters buried within the protein interior, and is generally in the $[Fe_4S_4]^{+2}$ state when poised above -530 mV (13, 14). Given its position at the center of the dimer, a structural role in stabilizing the dimer has been hypothesized, however recent mutational work suggests this is not the case (15). To date, no sign of the D Cluster has been observed in EPR spectra collected on CODH-II from *C. hydrogenoformans*,(2) but recent work in *Rhodospirillum Rubrum* yields a rhombic spectrum with g = 2.042, 1.933 and 1.880 in the range of -250 mV to -400 mV (15).

Buried ~13 Å beneath the D Cluster is the B cluster, a ferredoxin-like iron-sulfur cluster with a reduction potential of -440 mV which transfers electrons between the D cluster and the more deeply buried C cluster (16). The EPR spectra of the B cluster obtained for CODH-II are rhombic, with g = 2.04, 1.94 and 1.90, (2, 17) and match those obtained for the B Cluster of CODH from *Moorella thermoacetica* (16). These values are remarkably similar to those reported for the D Cluster RrCODH (15). However, EPR spectra collected on CODH-II to-date have shown major differences to those collected on RrCODH, particularly in the $C_{red2}$ state, indicating that further investigation is needed to understand the electronic structure of the C cluster form CODH-II.

The C cluster is buried approximately 12 Å below the B cluster, and is the catalytic heart of CODH. The structure of the C cluster differs greatly from the B and D clusters, having essentially been opened, and one of the Fe replaced by Ni (18). The displaced Fe has several names: Ferrous component II,(19) the pendant iron,(13, 20), the unique iron site $Fe_u$(21) or $Fe_1$ (12, 22). The $Fe_u$ remains joined to the cluster via a $\mu^3$-sulfide, where it gains additional coordination from C295, H261 and $H_xO^{x-2}$,(6) where x depends on the redox poise of the C cluster. The cuboidal core of the cluster, inclusive of the Ni is anchored to the protein through four additional covalent thiolate bonds provided by C333, C446, C476 and C526, where C526 is bound to Ni which is bound to the [Fe3S4] core of the C cluster via two additional $\mu^3$-sulfides, forming an overall T geometry. Unlike the D and B clusters, the C cluster can adopt one of four redox states, each corresponding to a stage in its catalytic cycle (Figure 1, Figure S1) $C_{ox}$ is EPR silent (S = 0), and represents the fully oxidized inactive state, characterized by N(II) and a diamagnetic $[Fe_3S_4+Fe_u]^{+2}$ component. The one electron reduced species, $C_{red1}$ is S = 1/2 (E° ~ -100mV),(2) with EPR, Ni L-edge XAS and Mössbauer data indicating the Ni remains diamagnetic Ni(II), while the $[Fe_3S_4+Fe_u]$ component resembles a high-spin $[Fe_3S_4+Fe_u]^{+1}$ cluster with $Fe_u$ coupled to the rest of the cluster to yield the final half-integer spin system (16, 23, 24). These measurements were taken on CODH from *Rhodospirillum rubrum* and *M. thermoacetica*. A further one-electron reduction yields the S = 0 $C_{int}$ species with Mössbauer spectra typical of a $[Fe_4S_4]$ cluster, although the B and D clusters contribute to the signal as well (24). A final single electron reduction yields $C_{red2}$, another S = ½ state with g values of 1.99, 1.87 and 1.74,(2) in general agreement with $C_{red2}$ determined from *M. thermoacetica* (16). The potential of the $C_{red1}/C_{red2}$ couple is -520 to -530 mV,(16, 25) matching with the potential of the $CO_2$/CO couple, which ranges from -499 mV at pH 6 to -617 mV at pH 8 (26). The assignment of oxidation states for $C_{red2}$ has been contentious, with Ni proposed to be in a zero valent state, or as a Ni(II)-hydride species, (27–29) and the formal assignment of $C_{red1}/C_{red2}$ as Ni(II)/Ni(0) was the recent focus of extensive computational work focusing on the binding and release of $CO_2$ to and from the C cluster (21).

Substantial work has been invested in untangling the structure of CODH, and its active site, using crystallographic and spectroscopic approaches. These studies have greatly advanced our understanding of the structure of CODH, and potential binding modes for substrates and substrate mimics (6, 12, 19, 28, 30) They have also sparked debate over the structure of the active site, including the binding mode of CO/$CN^-$ and the presence of a bridging $S^{2-}$ or $Cl^-$ between the Ni and $Fe_u$ (12, 22, 30, 31). However, even the crystal structures with the best resolution suffer from significant uncertainty in the structure of the C cluster, exhibiting multiple conformations for key atoms,(6, 12, 22, 28, 32) and different structures have been obtained from similar treatments, for example poising at -320 mV with DTT to recreate the $C_{red1}$ intermediate has resulted in structures with and without a bridging $S^{2-}$ or $Cl^{-}$ between the Ni and $Fe_u$ (22, 28). Recently, Basak et al. published a detailed crystallographic study investigating the $CO_2$ reduction by the C cluster (33). But this too suffers from inconsistencies with prior data, for example a $C_{red1}$ state (PDB: 9FPG) reduced with Ti(III) at pH 6.0 yielded a bridging -OH ligand between the Ni and $Fe_u$, inconsistent with prior $C_{red1}$ structures (1SU7, 3B53). This study further underscores the complexity of CODH, and investigations into its catalytic cycle. This is emphasized, notably in their reported structure (PDB: 9FPO) which claims to capture both CO bound and $CO_2$ bound species. As we detail here, there is a high degree of uncertainty, and even confusion in the existing crystallographic literature pertaining to the catalytic intermediates of CODH. Spectroscopic investigations using Ni K-edge XAS have not necessarily fared better, with existing data often showing little to no change among the various intermediates, suggesting a lack of conversion (34–37). Thus, there is still a need for a detailed structural description of the C cluster, and the Ni site in particular. In this study, we present detailed XAS and EPR data detailing the geometry and electronic structures of the C cluster in the $C_{ox}$, $C_{red1}$ and $C_{red2}$ states.

### Section III: Preliminary analysis of the Cred1 EPR at High Power

At 46 mW, CODH exhibits signal from the C cluster in two conformations, simulated with g = 1.63, 1.85 and 1.97 and the other with 1.74, 1.81 and 2.00, along with a small proportion of reduced B cluster and benzyl viologen. Precise quantification of each species is complicated by differences strain, but the two conformations appear in an approximate 3:1 ratio. All fit parameters and relative proportions presented in Table S2. Two EPR-detectable conformations for $C_{red1}$ in the same sample have not yet been reported. Values of g in the vicinity of 1.74 are typically diagnostic for the fully reduced $C_{red2}$ state (16, 25, 33), which does not accumulate in any appreciable quantity here due to the large (170 mV) difference in reduction potentials. For CODH-II $C_{red1}$, values of g = 1.73 and 1.64 have been reported previously (33, 38), but not on the same sample. Meanwhile, multiple EPR-detectable conformations have been reported for [Fe4S4] cluster systems (39–41), attributed to two dominant configurations. Recent, detailed work in the [Fe,Fe]-hydrogenase *Cr*HydA1 found that two EPR-detectable conformations of its $4Fe_H$ cluster were due to two dominant orientations of two positively charged second-sphere residues (42). Given the complexity of the C cluster and its surrounding residues, it is unclear at this moment whether key, positively charged second sphere residues like K563, or the protonation state of water bound to $Fe_u$ may be responsible, but it can be said that neither set of g values reported here match with those previously reported for Ni-deficient $C_{red1}$ (33). We plan to investigate these conformations with further mutational and large-scale computational studies.

### Section IV: A Brief history of substrate-free CODH-II Crystallography

CODH has a long history with crystallography, which has been a critical technique in advancing our understanding of the structure and mechanism of the enzyme. First solved by Dobbek et al in 2001(43) (PDB 1JJY), the C cluster was revealed to be a the distorted Ni-substituted iron sulfur cubane it is known to be today, with a $\mu_2$ sulfide bridging the Ni and $Fe_u$, yielding a square pyramidal Ni. We must note that, within 2 months of this structure, Drennan et al. (44) published their structure of RrCODH, which did not have this bridging sulfide. However, the sulfide was later reaffirmed by Dobbek et al. 2004, and PDB:1JJY was superseded in the PDB by 1SU8, a structure of CODH-II reduced by dithionite under and $N_2$ atmosphere, which showed the same bridging sulfide as the 2001 study. However, in their 2007 study, (28) Jeoung and Dobbek publish new structures of catalytically competent CODH-II lacking the bridging sulfide. The explanation offered for the discrepancy is contradictory, however, with the acknowledgement that the enzyme used in those prior studies, as well as dissolved crystals exhibiting the bridging sulfide had high (~14,000 units $mg^{-1}$) activity, while the enzyme used in the 2007 work had comparable activity, and no sign of the bridging sulfide. In fact, they find that the location of the sulfide is the location of $CO_2$ binding. This discrepancy leads to the contradictory conclusion that the bridging sulfide must be systematically absent in the catalytically competent enzyme, and that it may be chemically or reductively displaced to activate the enzyme (28). However, no S anomalous map is provided to substantiate the placement of S in the bridging position. Later, in 2009 Kung et al. (31) put forward compelling evidence from MtCODH/ACS, in comparison with the earlier data from Dobbek et al. that the bridging sulfide is not necessary, and is actually inconsistent with the mechanism as understood at that time. This is quickly corroborated by Jeoung and Dobbek, who demonstrate that the CO analog $CN^-$ binds to this location as well (30). At this stage, Kung et al.(31) suggested that the bridging sulfide might exist in an inactive pre-catalytic state. This recently found support from the Dobbek group, who demonstrated that, upon short (1 minute) exposure to $O_2$, the C cluster experiences and influx of sulfide, with one occupying the $\mu_2$ bridging position, and another forming a disulfide bond with Cys294, and two waters at 2.09 Å from the Ni. While not an exhaustive summary of the history of CODH through the lens of crystallography, we would be remiss not to discuss the recent structures by Basak et al (33). Briefly, nine new structures are presented, reduced with either Ti(III) or Eu(II)-DTPA, in various states of substrate binding. We explore this in more detail in the discussion below.

As highlighted briefly here, CODH is a difficult system to work in, and the progress towards understanding its underlying mechanism has benefited greatly from crystallography. However, as demonstrated via the multi-year saga of the $\mu_2$ bridging sulfide, the crystallography data are conflicted on the identities of the CODH-II intermediates. Here we detail these conflicts and point to where the spectroscopic data we provide can be used to clarify the identities of $C_{ox}$, $C_{red1}$ and $C_{red2}$. Prior to the recent pH jump crystallographic study by Basak et al.(33) 13 structures of WT monofunctional CODH-II from *C. hydrogenoformans* are available in the PDB. However, only five structures have no substrate bound ($C_{red2}$: 3B51, 1SU8; $C_{red1}$: 3B53 and 1SU7; $C_{ox}$ 1SUF). Of the new structures presented by Basak et al.(33) , only three have no substrate bound. Of those structures, two are stated to be in the $C_{int}$ state (9FPF and 9FPN), leaving only the claimed $C_{red1}$ state, 9FPG. Here we present an analysis of these structures, focused on three regions: Ni and its ligands, $Fe_u$ – whether it is "in" or "out" (Figure S1) and it's ligands, and any interactions between Cys294 and the cubane $\mu_3$ sulfides – as it is implicated in C cluster maturation (18), and finally we present a RSZD analysis of the C cluster in each structure. 1SUF is presented as the "inactive state" of CODH-II (22). The Ni is weakly bound by 2 O at 2.84 Å, but the Ni (and all Fe) show signs of over fitting, yielding Fo-Fc density peaks extending to a 31σ contour (Ni), 48σ contour (Fe2 and Fe3) and 20σ contour ($Fe_u$). This is likely an artifact of artificially low B-factors, which range from 0.08 to 7.64 for the metals, with all occupancies constrained to 100% (Table S5). Moreover, $Fe_u$ is refined to 30% occupancy in the "in" position with an implausibly close 1.97Å bond from the Ni and no evidence in the 2Fo-Fc map of His261 moving in to maintain coordination to the $Fe_{u,i}$ as proposed in a recent 2025 paper (33). Here, the authors propose that $Fe_u$ moves in between the "in" and "out" positions as a part of a dynamic process wherein $Fe_u$ moves between the two positions in response to the redox poise of the C cluster, the pH, and presence of CO while maintaining coordination by His261(33). The lack of His261 movement to coordinate $Fe_{u,I}$ in 1SUF results in an implausibly 3 coordinate $Fe_u$, gaining coordination by Cys526. It is also worth noting that, in the structures which have $Fe_{u,I}$, it comes improbably close to the Ni (~2.4 Å in 9FPG, 3B51 and 3B53), suggesting that the Ni is absent in this configuration of $Fe_u$, and it is thus a sign of cluster degradation.

Structures of $C_{red1}$ do not fare much better. In addition to 1SU7 and 3B53, 9FPG was recently deposited. 1SU7 was formed via incubation of CODH-II crystals with 2 mM DTT under a nitrogen atmosphere at pH 7.5. The resulting structure has a bridging $\mu_3$ sulfide, and it is now known that this is not a component of the active enzyme (28, 31). Apart from the sulfide, the Ni is bound to one water at 2.29 Å with a high b-factor of 34.8 Å$^2$ (compared to Ni's 12.45 Å$^2$) owing to having been refined to a position just at the edge of the 2Fo-Fc difference density. Moreover, there is at least one additional unexplained portion of electron density above the Ni which may be due to additional waters. Cys526 displays negative Fo-Fc difference density, with a positive density peak suggesting an alternate conformation away from the Ni atom, and is likely indicative of the location of Cys526 when Ni is absent. The occupancies of the C cluster atoms are all above 0.9, despite an Fe Fo-Fc density peak remaining at 10σ contour, and at 13σ contour for Ni. While $Fe_u$ is only in the out conformation, Fe2 displays two conformations with one binding to Cys294 with an occupancy of 30% (vs 100% for Cys294). This bond is not observed in the other structures and is a signal of cluster degradation as Fe2 as this second position is 1.66 Å removed from its position in the cubane. 3B53 improves on 1SU7 and was made via incubation of 7 mM DTT at pH 8 (28). The Ni shows no sign of a bound water, and the $Fe_u$ has 60% occupancy in the out position with 30% in the in position. However, the $Fe_{u,i}$ loses His261 coordination, and is assigned to bind with the conformation of Cys295 which has 70% occupancy. 9FPG is the most recent $C_{red1}$ structure, made via incubation of the CODH crystal in Ti(III)-EDTA at pH 6 (33). No water is bound to the apical position of Ni, as in 1SU7, but substantial unmodelled density is present in that position. The Ni is bridged to the $Fe_u$ via an OH, which was not observed in prior structures. The OH is described as bound to $Fe_u$ in the out position, but was refined as though bound to both $Fe_u$ positions. The occupancy of the metals exceeds that of the sulfides, with Fe density in the cubane with a persisting difference density peak extending to a 5.5 σ contour. Cys294 is found in two locations, with the B conformation having 15% occupancy and forming a 1.94 Å bond to the cubane S3 sulfide, which may indicate some damage to the structure, as similar Cys-sulfide interactions are implicated in both cluster maturation and degradation (12, 18).

Structures of $C_{red2}$ are comparable. 1SU8 has been discussed previously,(31) but our critiques are summarized here. 1SU8 was made via incubation with dithionite at pH 7.5. The Ni is bound to a water in the apical position, but the water is misplaced, resulting in a difference density peak at 3σ contour. The Ni is overfit, based on Fo-Fc density peak up to 13.3σ contour. 3B51 was made via incubation with Ti(III) citrate at pH 8.0, and shows a Ni with no water bound. $Fe_u$ is present in both the in and out conformation, with a water bound to the $Fe_u$ in the out position, and no His261 coordination on the $Fe_{u,i}$.

We have kept this analysis of the crystal literature to those structures with no substrate bound, but recent substrate bound structures fair no better. For example, the CO bound structure 9FPO was described as Eu(II)-reduced, pH 6.0 + CO crystal, $C_{int,o}$ + CO state (33). Here, there are two conformations of Ni, the majority Nia (50% Occ.) and Nib (24% Occ), with water (15% Occ.) and CO (50% Occ.) bound to Nib and Nia, respectively, with a long 2.27 Å Ni-C distance. $CO_2$ is also present (26% Occ.), bound to $Fe_{u,o}$ (55% Occ.) while a water (30% Occ.) is bound to the $Fe_{u,I}$ (30% Occ.) in the same position as $CO_2$. $CO_2$ is also designated in the structure as bound to Nia, a clear indication that multiple species are present in the same crystal structure, but which are not fully distinguishable from each other. The authors indicate that the modelling approach taken was a region-by-region approach, as opposed to fitting full CO and $CO_2$ bound states, as it is implausible that Nia would be bound to CO and $CO_2$ simultaneously. Our understanding of the C cluster is ongoing. The XAS compliments the existing conflicted crystal data, resolving the coordination structure of Ni in each state.

### Section V: Comprehensive RSZD Survey of All Available CODH Structures

Modelling the crystal structure of CODH is a difficult task. CODH is plagued by several factors that make analyzing metalloproteins difficult such as metal loading in sample preparation, oxygen sensitivity and radiation damage (45). These three factors can result in crystallography data of different species all being present in one crystal structure. Modelling such an ensemble may lead to models with uncertain bond lengths and impractical occupancies that are often less than 1 and, in most cases, are allowed to be refined by individual atom due to C-cluster degradation. This has resulted in high uncertainty in the C cluster region in crystallographic studies of CODH, with a median RSZD score of 6.6 σ across all available structures. This bolsters the need for techniques such as XAS to verify crystallography data. For example, in CODH of *Desulfovibrio vulgaris* (PDB:6B6W) solved at 1.72 Å resolution with two alternate conformations in which one includes and one excludes the Ni atom with RSZD scores of the C-cluster at 4.3 σ. Even high-resolution structures are difficult to deconvolute. A recent structure of CODH from *C. hydrogenoformans* in the $C_{red2}$ state bound with $CO_2$ solved at 1.01 resolution (PDB: 9FPH) includes difference density peaks that result in an RSZD score of the stand-alone C-cluster of 12.5 σ. The median of RSZD scores of all CODH crystal structures with an intact C-cluster have been found to be 6.1 σ (Table S12), demonstrating that CODH's C cluster is challenging to model accurately to the data collected. Furthermore, structures that have a low RSZD score such as CODH from *Rhodospirillum Rubrum* (PDB: 9GYB) solved at 2.90 Å contains a C cluster RSZD score of 1.1, but the cluster has unusually high B-factors to compensate. The RSZD scores for all available CODH structures have been calculated, and are presented in Table S12.

### Section VI: Constructive and destructive Interference of Ni-Fe paths in EXAFS of the C cluster

The effects of destructive interference of competing Fe-Fe paths has been well documented in iron sulfur cubanes (46, 47). Successvive reduction is generally associated increased structural disorder (variation in Me-Me distances), which can manifest as a larger Debye-Waller factor ($\sigma^2$) for that interaction in the EXAFS (46–49). The Debye waller factor has both static and vibrational component, the later of which can be minimizxed by measurement at low temperature. Larger values of $\sigma^2$ reflecting greater disorder effectively attenuate the EXAFS signal for the associated metal-metal scattering interaction (Figure S11). Alternatively, the EXAFS signal can be attenuated via destrcutive interfence of competing metal-metal paths. Typically, an offset of 0.2 Å between two metal-metal scattering distances will result in substantial attenuation of the signal, although it may leave a unique beating pattern which can be used to distinguish the two interactions. Shown in Figure S11 are competing Ni-Fe paths, and their scattering sum illustrating the sensitivity of EXAFS towards small changes in the distance.

Clearly, this signal can be attenuated to the noise level (i.e. no obvious metal-metal contributions beyond the first shell), straining good data practice if one attempts to model the interaction in the fit. However, this appears to be where the C lcuster departs in behavior from conventional iron sulfur cubanes. In the absence of substrate, those Ni-Fe contributions gain intenisty as the C cluster is reduced to $C_{red1}$ and then $C_{red2}$. Models of the Ni-Fe interactions demonstrate that this is due to both decrease in structural disorder ($\sigma^2$ decreases) as well as a decrease in destructiuve interference, as the difference in the modelled Ni-Fe lengths decreases from 0.23 Å ($C_{ox}$) to 0.18 Å ($C_{red2}$) (Figure S14). On the binding of $CN^-$ or $OCN^-$ to $C_{red1}$ and $C_{red2}$ (respectivley), the Ni-Fe contribution gains incredible prominance, relative to their substrate-free forms. This manifests as both much smaller $\sigma^2$ and smaller differences in the two Ni-Fe lengths, which each contribute to the larger Ni-Fe signal. In both species, substrate binding reulst in a reorganization of the cluster. In $C_{red1}$, $CN^-$ binding leads to a 0.04 Å expansion of the first Ni-Fe path, and a 0.03 Å contraction of the second, and an overall 0.07 contraction in the distance between the two Fe paths, greatly reducing the magnitude of destructive interference. In $C_{red2}$, $OCN^-$ binding elongates the Ni-Fe paths by a substantial 0.18 Å and 0.17 Å, suggesting expansion of the cluster as the ligand is accomodated. Given that the substrate bound and free samples were generated form the same protocol and batch of CODH-II, having such an obvious Fe signal in the substrate bound forms guides the analysis of the subtsrate free forms, and led to the demonstration of how the Ni-Fe contribution is attenuated in the $C_{ox}$.

### Figures S1 to S15

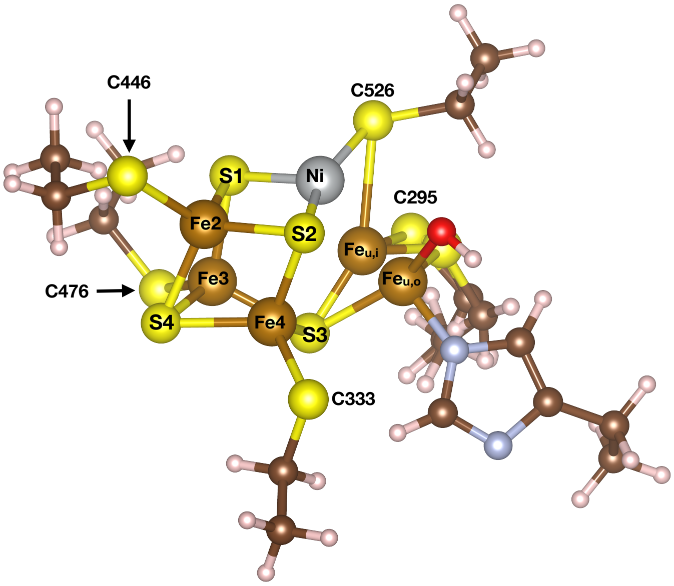

**Figure S1.** Numbering scheme used for atoms constituting the C cluster in this work. The unique Fe is shown in the in (Feu,i) and out (Feu,o) conformations described by Basak et al. 2025.

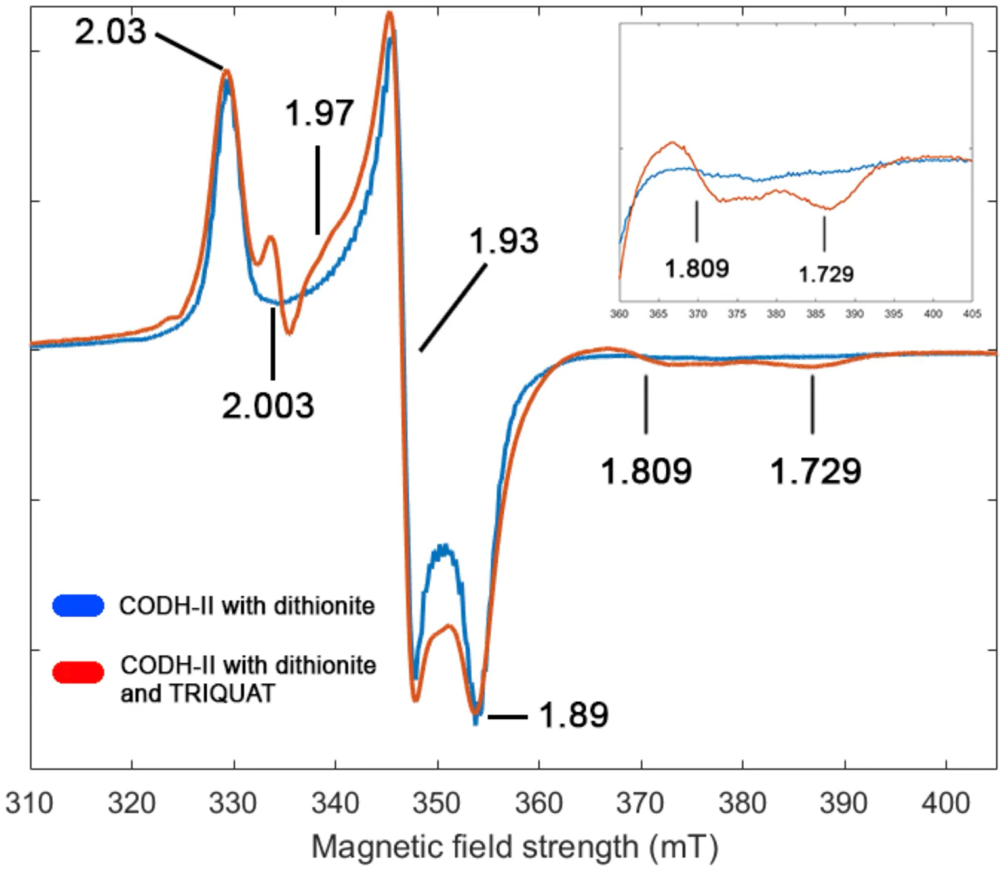

**Figure S2.** X-band EPR spectra of CODH-II treated with either dithionite only (blue) or dithionite and TRIQUAT (red). All protein concentrations were 250 uM as determined by Rose-Bengal assay. Dithionite only sample was prepared with 5 mM dithionite. The dithionite and TRIQUAT sample was prepared with 5 mM dithionite and 500 uM TRIQUAT. TRIQUAT is responsible for the g = 2.0 feature. TRIQUAT. Microwave frequency: 9.376 GHz. Modulation amplitude: 10 G. Conversion time: 163.8 s; time constant: 81.92 ms. Microwave power: 2 mW; attenuation: 20 dB. Receiver antenna gain: 5.03x 10^3.

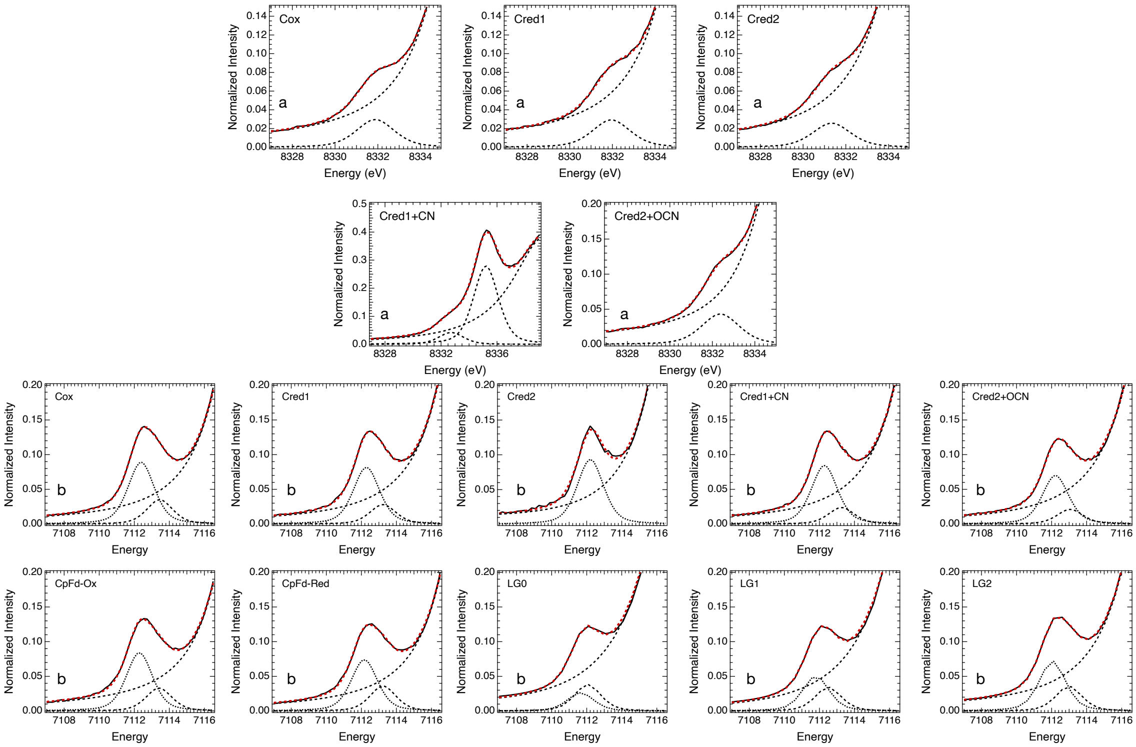

**Figure S3.** Pseudo-Voigt peak deconvolution of the (a) Ni K-pre-edge and (b) Fe K-pre-edge. Samples LG0-LG2 the [Fe4S4]0 thru [Fe4S4]2+ and are from https://doi.org/10.1021/jacsau.4c00213 and have been made available for use by the authors under license at https://doi.org/10.3929/ethz-b-000742321.

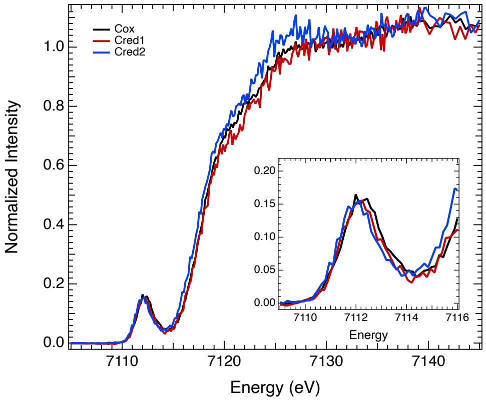

**Figure S4.** Fe Kα-HERFD for the three CODH intermediates under consideration. (*inset*) The expanded K-pre-edge region highlighting the 1s→3d transition.

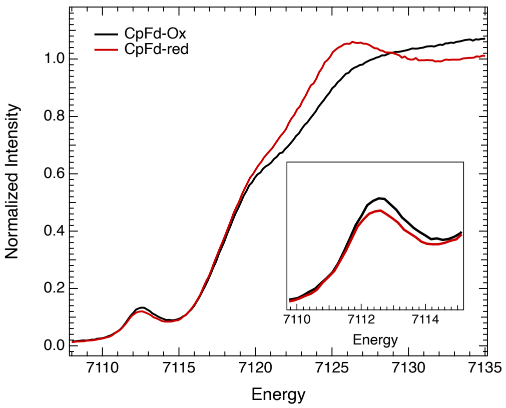

**Figure S5.** Fe K-edge XAS. (Left) Oxidized and reduced CpFd (courtesy of G. Graham).

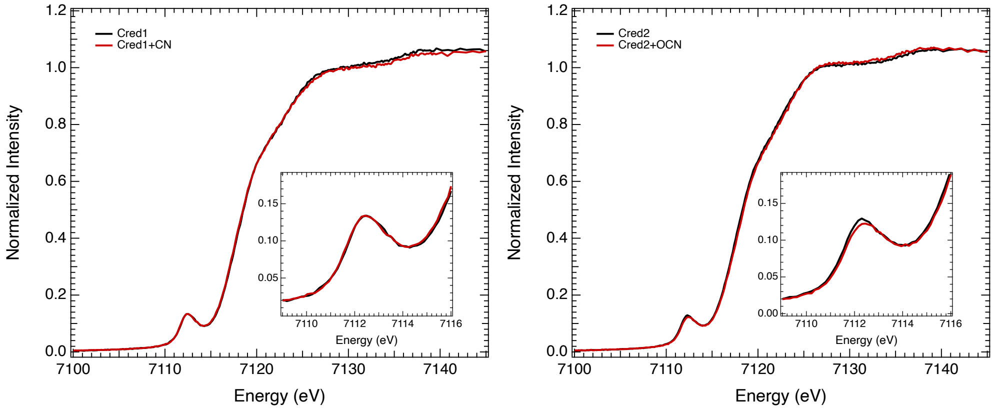

**Figure S6.** Fe K-edge XAS. (Left) $C_{red1}$ (Right) $C_{red2}$

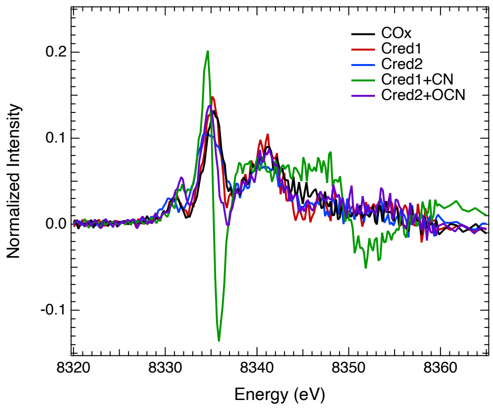

**Figure S7.** First derivatives of the Ni K-edge XAS data.

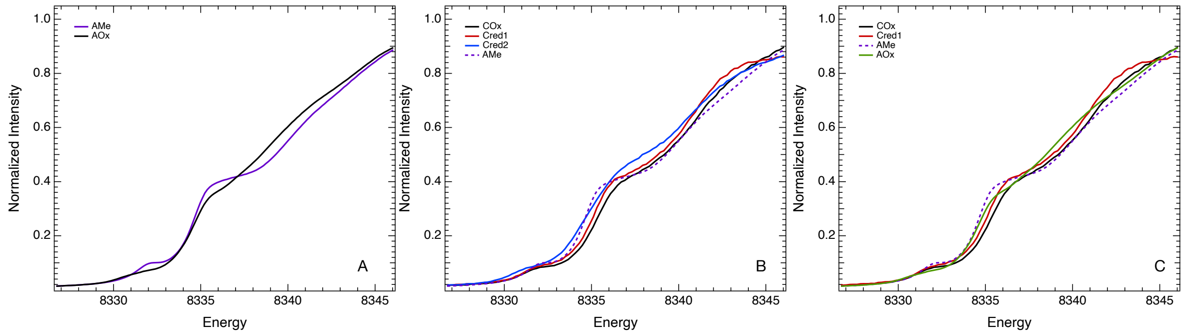

**Figure S8.** Ni K-edge comparison of the A cluster from ACS (*N. thermoacetica*) with CODH-II. (A) The Ni K-edge spectra of oxidized ($A_{Ox}$) and methylated ($A_{Me}$). (B) Ni K-edge comparison of the planar $A_{Me}$ with the three CODH intermediates, highlighting the difference in the riding edge feature at 8336.3 eV. (C) The comparison of the two ACS intermediates with $C_{ox}$ and $C_{red1}$, which have the sharpest rising edges of the three intermediates.

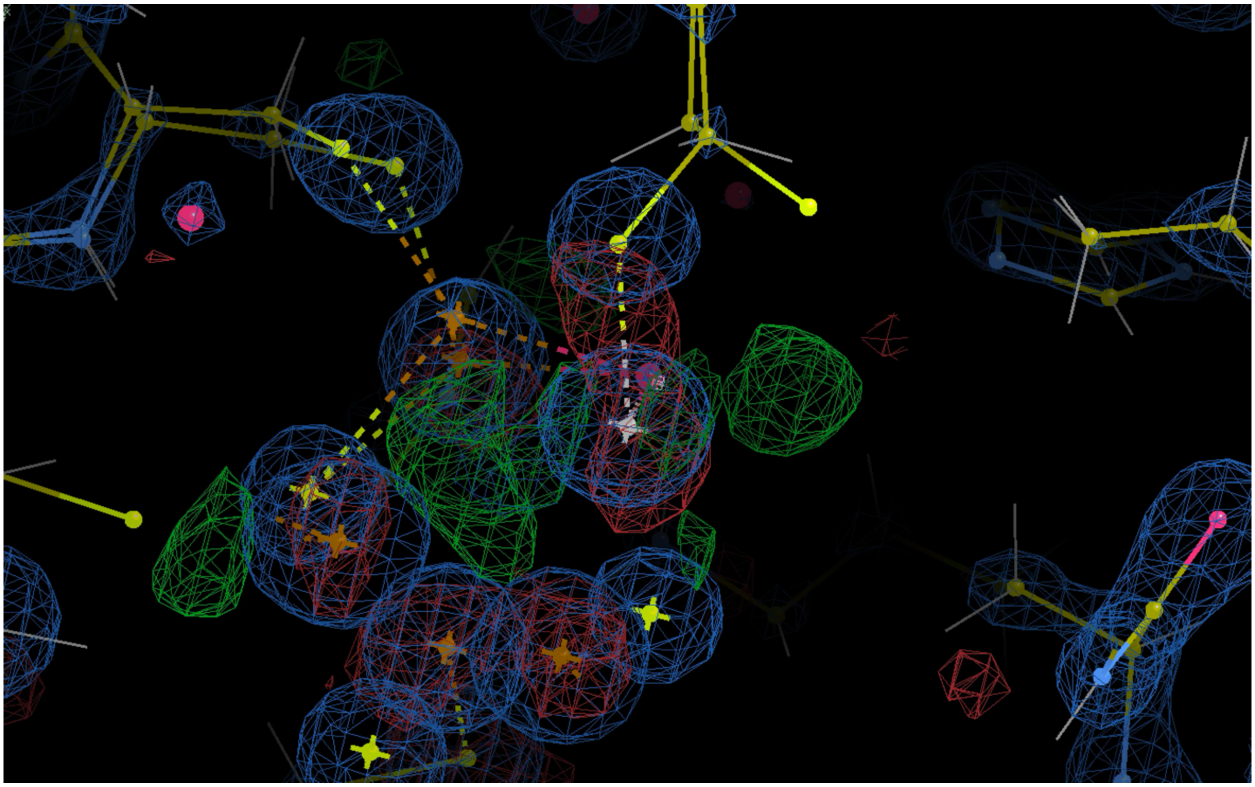

**Figure S9.** Density map of 9FPG showing overfit between Cys526 and Ni contoured to 3σ. 2Fo-Fc (blue), Fo-Fc showing areas of overfit (red) and under fit (green).

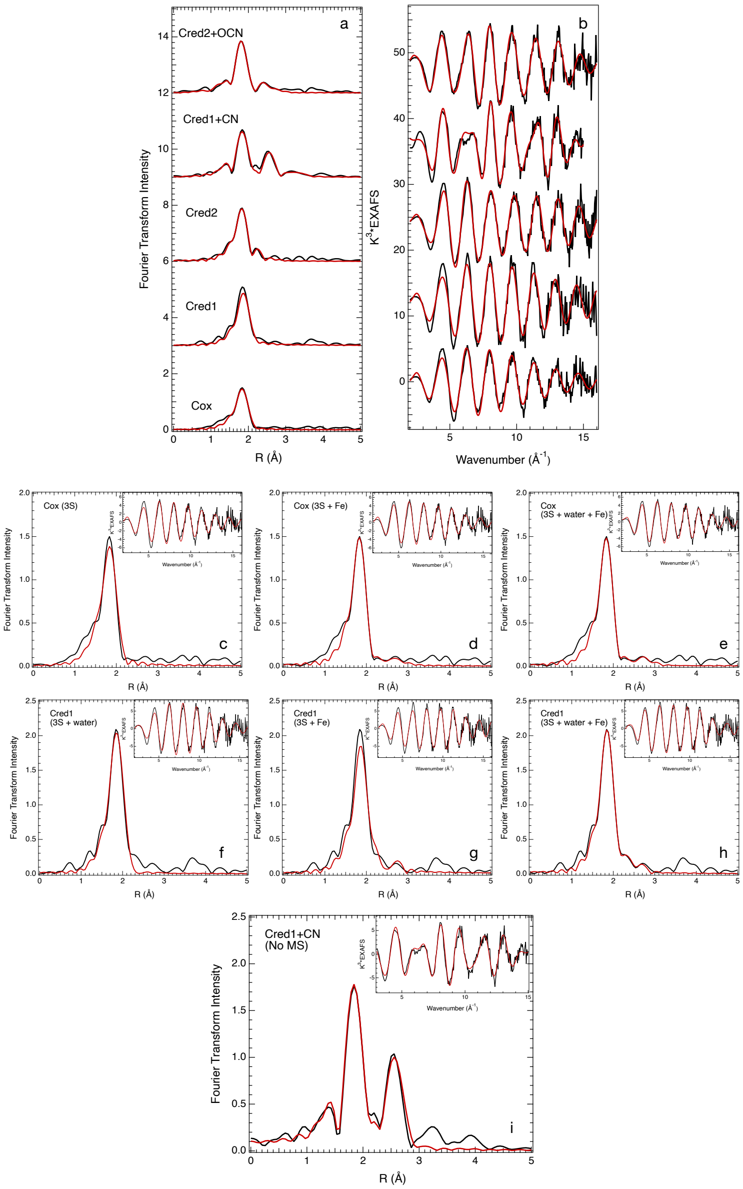

**Figure S10.** Ni K-edge EXAFS fits. (a-b) Minimal fits for all samples. Corresponding fit parameters are presented in Table 1. (c-d) Alternative fits for Cox. c) a single S shell is insufficient to model the Ni-S interaction. d) Two Fe shells are modeled with a split S shell. e) A single S shell is sufficient to model the Ni-S interaction when a water at 2.09 Å is included. However, this is compensated for by a ~40% increase in the $\sigma^2$ of the Ni-S shell, relative to its value in fit (c), and a 3x increase relative to the minimal fit. The Ni-Fe paths remain relatively unperturbed whether a water or split S shell is used. (f-h) Alternative fits for Cred1. f) The minimal fit presented in (a) is enhanced by the addition of two waters at 2.41 Å. g) A single Ni-S shell with Ni-Fe contributions added. The Fe contributions are insufficient to enhance the fit of the Ni-S shell. h) The same as g), but with two waters modeled at 2.43 Å to enhance the fitting of the first shell. The position of the two Fe remains relatively unperturbed whether a water is included in the first shell or not. i) An alternative fit for Cred1+CN where the multiple scattering is excluded, revealing a comparable fit quality. The position of the Fe remains relatively unperturbed regardless of whether the multiple scattering is included. Corresponding fit parameters are available in Table S5.

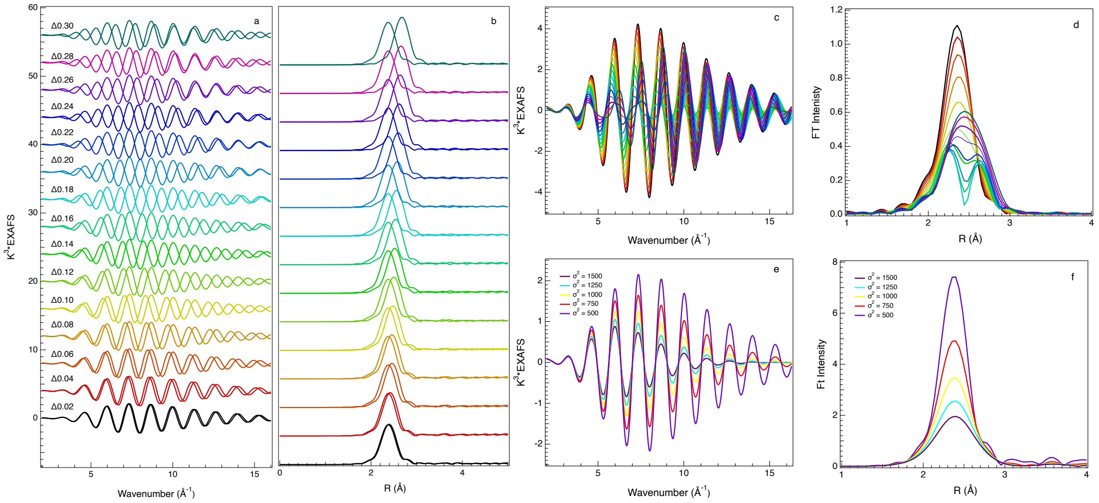

**Figure S11.** Effects of attenuation on the Ni-Fe EXAFS signal. A) the signal from 2 Ni-Fe paths, one anchored at 2.65 Å, and the second moved further in units of 0.02 Å out to 2.95 Å. B) The Fourier-transforms of the individual two paths from (a). C) The sums of the two paths from (a) highlighting how destructive interference manifests as a function of path separation in the $K^3$-weighted EXAFS. D) The Fourier-transform of the sum of the paths presented in (c), illustrating how the destructive interference manifests in R-space. Note that the colors have been kept consistent from panels A-D. E) The effect of increasing the Debye-Waller term on the EXAFS of an Ni-Fe path set at 2.65 Å. The magnitude of the Debye-Waller term was varied between 500 and 1500 (all values multiplied by $10^{-4}$). F) The Fourier-transform of the paths from (e) illustrating how an increasing Debye-Waller factor dampens the Ni-Fe EXAFS contribution in R-space.

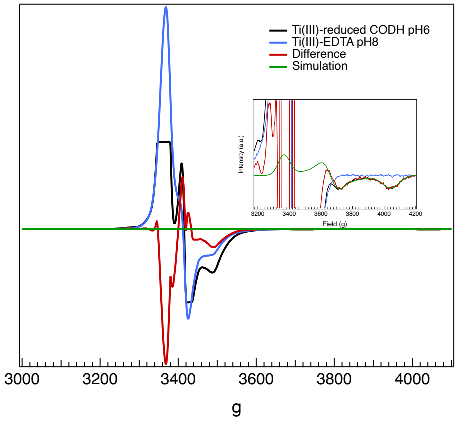

**Figure S12.** Assessment of prior Cred1 EPR from Basak et al. 2025. Subtraction of Ti(III)-EDTA (multiplied by 40) from CODH reduced by Ti(III)-EDTA at pH 6.0 to recreate simulation (in green). The Ti(III)-EDTA signal was not of equal intensity in both samples, being less intense in the Ti(III)-EDTA control requiring multiplication such that the resulting subtracted spectrum reproduced the intensity of the g = 1.82 and 1.64 features (*inset*). Subtracted spectrum highlights that obtaining the g = 1.97 is not possible due to cut-off in the CODH+Ti(III) spectrum, characteristic of detector diode saturation.

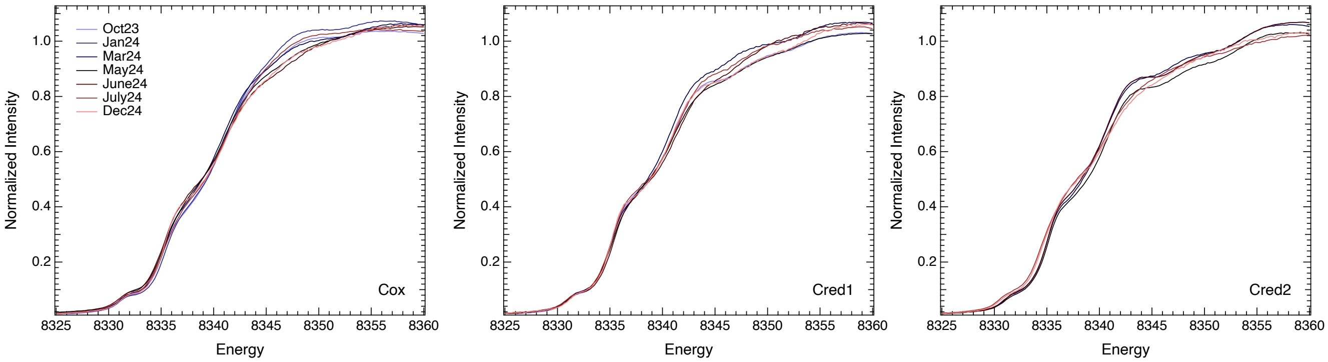

**Figure S13.** Iterations of preparation protocols for CODH intermediates highlighting variability in absence of the protocol detailed in this study.

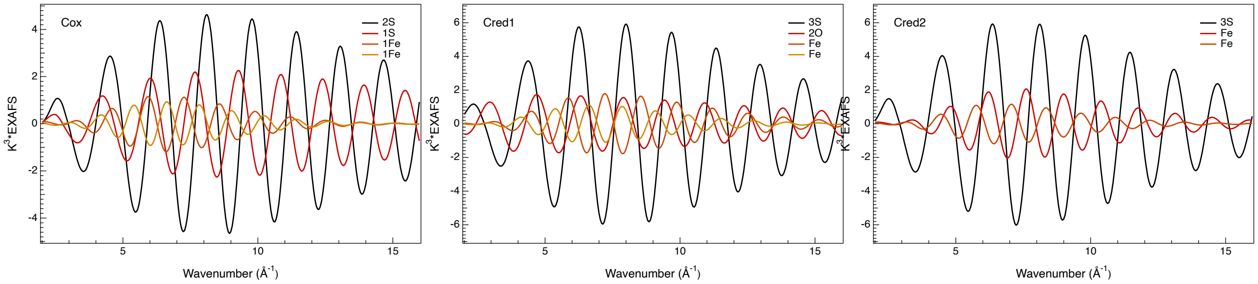

**Figure S14.** Contributions of each shell to the best-fit EXAFS (iron paths included) for Cox, Cred1 and Cred2, corresponding to fits S10d, S10h and S10a, respectively.

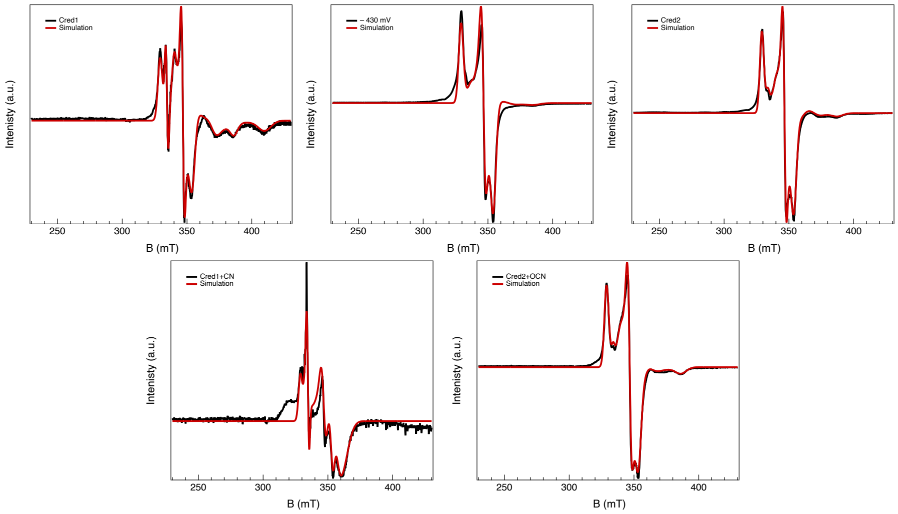

**Figure S15.** Simulations of the EPR data collected at –370 mV (Cred1), –430 mV (B Cluster), –540 mV (Cred2), –370 mV in the presence of $CN^{-}$ (Cred1+CN) and -540 mV in the presence of $OCN^{-}$ (Cred2+OCN). Simulation parameters are presented in Table S2.

### Tables S1 to S12

**Table S1. PDB Survey of all available CODH structures for Ni-Fe and Ni-S distances**

| PDB Entry Alternate Location | Ni_Feu1_distance | Ni_Feu2_distance | Ni_Fe2_distance | Ni_S1_distance | Ni_S2_distance | Ni_S3_distance |
|---|---|---|---|---|---|---|
| 9gyb_NI_multi_hetero_d7.00_cluster1.xyz | 2.581 | | 2.436 | 2.291 | 2.293 | 2.944 |
| 8cjb_NI_multi_hetero_d7.00_cluster1_altlocLIGA.xyz | 3.034 | | 2.538 | 1.969 | 2.278 | 2.323 |
| 6vwy_NI_multi_hetero_d7.00_cluster1_altlocNiA.xyz | 2.411 | | 2.541 | 2.221 | 2.258 | 2.412 |
| 9gyb_NI_multi_hetero_d7.00_cluster5.xyz | 2.612 | | 2.565 | 1.971 | 2.287 | 2.289 |
| 6vwy_NI_multi_hetero_d7.00_cluster2_altlocNiA.xyz | 2.464 | | 2.574 | 2.205 | 2.255 | 2.432 |
| 9fpn_NI_multi_hetero_d7.00_cluster1_altlocLIGA.xyz | 2.6 | | 2.575 | 2.111 | 2.125 | 2.177 |
| 1jqk_NI_multi_hetero_d7.00_cluster5.xyz | 2.607 | | 2.608 | 2.152 | 2.224 | 2.244 |
| 1jqk_NI_multi_hetero_d7.00_cluster3.xyz | 2.64 | | 2.608 | 2.188 | 2.206 | 2.236 |
| 3b51_NI_multi_hetero_d7.00_cluster1_altlocLIGA.xyz | 2.442 | | 2.618 | 2.028 | 2.161 | 2.169 |
| 3b51_NI_multi_hetero_d7.00_cluster1_altlocLIGB.xyz | 2.875 | | 2.618 | 2.028 | 2.161 | 2.169 |
| 1jqk_NI_multi_hetero_d7.00_cluster4.xyz | 2.682 | | 2.635 | 2.183 | 2.195 | 2.251 |
| 9gyb_NI_multi_hetero_d7.00_cluster4.xyz | 2.601 | | 2.64 | 1.972 | 2.3 | 2.3 |
| 9fnj_NI_multi_hetero_d7.00_cluster1.xyz | 2.534 | 3.674 | 2.643 | 2.164 | 2.301 | 2.812 |
| 3b53_NI_multi_hetero_d7.00_cluster1_altlocLIGA.xyz | 2.383 | | 2.649 | 2.071 | 2.137 | 2.17 |
| 3b53_NI_multi_hetero_d7.00_cluster1_altlocLIGB.xyz | 2.854 | | 2.649 | 2.071 | 2.137 | 2.17 |
| 9c0t_NI_multi_hetero_d7.00_cluster1.xyz | 3.169 | | 2.649 | 2.249 | 2.32 | 2.335 |
| 1jqk_NI_multi_hetero_d7.00_cluster1.xyz | 2.616 | | 2.651 | 2.206 | 2.216 | 2.265 |
| 7zkk_NI_multi_hetero_d7.00_cluster1_altlocLIGA.xyz | 2.834 | | 2.652 | 2.065 | 2.251 | 2.341 |
| 6b6v_NI_multi_hetero_d7.00_cluster1.xyz | 2.579 | | 2.661 | 2.181 | 2.223 | 2.306 |
| 6yu9_NI_multi_hetero_d7.00_cluster1.xyz | 2.793 | | 2.662 | 2.236 | 2.299 | 2.52 |
| 6yu9_NI_multi_hetero_d7.00_cluster4.xyz | 2.78 | | 2.665 | 2.279 | 2.371 | 2.392 |
| 1jqk_NI_multi_hetero_d7.00_cluster6.xyz | 2.543 | | 2.667 | 2.202 | 2.256 | 2.259 |
| 7zxj_NI_multi_hetero_d7.00_cluster1_altlocLIGA.xyz | 2.679 | | 2.667 | 2.047 | 2.077 | 2.252 |
| 7zxj_NI_multi_hetero_d7.00_cluster1_altlocLIGC.xyz | | | 2.667 | 2.047 | 2.077 | 2.252 |
| 7zkj_NI_multi_hetero_d7.00_cluster1_altlocLIGA.xyz | 2.882 | | 2.675 | 2.193 | 2.222 | 2.306 |
| 6b6x_NI_multi_hetero_d7.00_cluster1_altlocLIGA.xyz | 2.999 | | 2.676 | 2.157 | 2.272 | 2.277 |
| 1oao_NI_multi_hetero_d7.00_cluster1_altlocLIGB.xyz | 3.554 | 3.579 | 2.676 | 2.29 | 2.291 | 2.303 |
| 6elq_NI_multi_hetero_d7.00_cluster3.xyz | 3.148 | | 2.679 | 1.96 | 2.051 | 2.414 |
| 6elq_NI_multi_hetero_d7.00_cluster1.xyz | 3.17 | | 2.679 | 2.017 | 2.053 | 2.413 |
| 9fpf_NI_multi_hetero_d7.00_cluster1_altlocLIGA.xyz | 2.878 | | 2.68 | 2.104 | 2.155 | 2.225 |
| 6x5k_NI_multi_hetero_d7.00_cluster2_altlocLIGA.xyz | 2.964 | | 2.682 | 2.181 | 2.314 | 2.345 |
| 7zxc_NI_multi_hetero_d7.00_cluster1_altlocLIGA.xyz | 2.3 | | 2.687 | 1.955 | 2.144 | 2.311 |
| 6b6v_NI_multi_hetero_d7.00_cluster2.xyz | 2.631 | | 2.687 | 2.163 | 2.241 | 2.308 |
| 7zxc_NI_multi_hetero_d7.00_cluster1_altlocLIGB.xyz | 2.789 | | 2.687 | 1.955 | 2.144 | 2.311 |
| 1oao_NI_multi_hetero_d7.00_cluster2_altlocLIGB.xyz | 3.487 | 3.63 | 2.687 | 2.311 | 2.325 | 2.361 |
| 3cf4_NI_multi_hetero_d7.00_cluster1.xyz | 3.158 | | 2.696 | 2.232 | 2.269 | 2.316 |
| 1jqk_NI_multi_hetero_d7.00_cluster2.xyz | 2.538 | | 2.713 | 2.223 | 2.255 | 2.312 |
| 3i04_NI_multi_hetero_d7.00_cluster4.xyz | 3.252 | | 2.717 | 2.236 | 2.311 | 2.317 |
| 3i01_NI_multi_hetero_d7.00_cluster1.xyz | 2.845 | | 2.721 | 2.156 | 2.219 | 2.244 |
| 9c0q_NI_multi_hetero_d7.00_cluster1.xyz | 2.421 | | 2.726 | 2.229 | 2.315 | 2.341 |
| 9c0q_NI_multi_hetero_d7.00_cluster2.xyz | 2.432 | | 2.728 | 2.23 | 2.315 | 2.34 |
| 6b6x_NI_multi_hetero_d7.00_cluster2_altlocLIGA.xyz | 3.055 | | 2.728 | 2.149 | 2.308 | 2.309 |
| 3i01_NI_multi_hetero_d7.00_cluster2.xyz | 2.865 | | 2.732 | 2.136 | 2.238 | 2.239 |
| 6x5k_NI_multi_hetero_d7.00_cluster4.xyz | 3.099 | | 2.732 | 2.187 | 2.308 | 2.345 |
| 3i01_NI_multi_hetero_d7.00_cluster4.xyz | 2.904 | | 2.738 | 2.281 | 2.312 | 2.326 |
| 7zkv_NI_multi_hetero_d7.00_cluster1_altlocLIGA.xyz | 2.85 | | 2.739 | 2.097 | 2.219 | 2.328 |
| 6x5k_NI_multi_hetero_d7.00_cluster1_altlocLIGA.xyz | 2.87 | | 2.741 | 2.161 | 2.338 | 2.342 |
| 9c0s_NI_multi_hetero_d7.00_cluster1.xyz | 2.532 | | 2.743 | 2.337 | 2.342 | 2.354 |
| 9c0s_NI_multi_hetero_d7.00_cluster2.xyz | 2.507 | | 2.749 | 2.34 | 2.349 | 2.422 |
| 7zx5_NI_multi_hetero_d7.00_cluster1_altlocLIGA.xyz | 2.448 | | 2.758 | 2.076 | 2.158 | 2.163 |
| 9c0t_NI_multi_hetero_d7.00_cluster2.xyz | 2.517 | | 2.758 | 2.335 | 2.338 | 2.378 |
| 7zxl_NI_multi_hetero_d7.00_cluster1_altlocLIGA.xyz | 2.699 | | 2.759 | 1.953 | 2.03 | 2.323 |
| 7zxl_NI_multi_hetero_d7.00_cluster1_altlocLIGC.xyz | | | 2.759 | 1.953 | 2.03 | 2.323 |
| 7zxl_NI_multi_hetero_d7.00_cluster1_altlocLIGD.xyz | | | 2.759 | 1.953 | 2.03 | 2.323 |
| 7zx6_NI_multi_hetero_d7.00_cluster1_altlocLIGA.xyz | 2.128 | | 2.761 | 2.15 | 2.222 | 2.275 |
| 9c0r_NI_multi_hetero_d7.00_cluster1.xyz | 2.58 | | 2.77 | 2.319 | 2.331 | 2.34 |
| 8x9g_NI_multi_hetero_d7.00_cluster1.xyz | 3.228 | | 2.771 | 2.186 | 2.453 | 2.47 |
| 9c0r_NI_multi_hetero_d7.00_cluster2.xyz | 2.57 | | 2.774 | 2.318 | 2.33 | 2.342 |
| 6yu9_NI_multi_hetero_d7.00_cluster2.xyz | 2.705 | | 2.775 | 2.25 | 2.279 | 2.487 |
| 3i01_NI_multi_hetero_d7.00_cluster3.xyz | 2.83 | | 2.779 | 2.114 | 2.117 | 2.417 |
| 8x9e_NI_multi_hetero_d7.00_cluster1_altlocLIGA.xyz | 2.576 | | 2.783 | 2.16 | 2.424 | 2.664 |
| 8x9e_NI_multi_hetero_d7.00_cluster1_altlocLIGB.xyz | 2.576 | | 2.783 | 2.16 | 2.424 | 2.664 |
| 3b52_NI_multi_hetero_d7.00_cluster1_altlocLIGA.xyz | 2.408 | | 2.793 | 2.091 | 2.104 | 2.228 |
| 3b52_NI_multi_hetero_d7.00_cluster1_altlocLIGB.xyz | 2.756 | | 2.793 | 2.091 | 2.104 | 2.228 |
| 9fnc_NI_multi_hetero_d7.00_cluster1.xyz | 2.597 | | 2.794 | 2.11 | 2.179 | 2.3 |
| 3i04_NI_multi_hetero_d7.00_cluster2.xyz | 3.157 | | 2.794 | 2.206 | 2.206 | 2.351 |
| 1mjg_NI_multi_hetero_d7.00_cluster1.xyz | 2.668 | | 2.795 | 2.161 | 2.275 | 2.288 |
| 1mjg_NI_multi_hetero_d7.00_cluster2.xyz | 2.649 | | 2.8 | 2.162 | 2.271 | 2.291 |
| 9fub_NI_multi_hetero_d7.00_cluster1_altlocLIGA.xyz | 2.558 | | 2.804 | 2.111 | 2.17 | 2.32 |
| 9fub_NI_multi_hetero_d7.00_cluster1_altlocLIGB.xyz | 2.558 | | 2.804 | 2.111 | 2.17 | 2.32 |
| 9fu9_NI_multi_hetero_d7.00_cluster1_altlocLIGA.xyz | 2.658 | | 2.804 | 2.11 | 2.174 | 2.299 |
| 9fu9_NI_multi_hetero_d7.00_cluster1_altlocLIGB.xyz | 2.658 | | 2.804 | 2.11 | 2.174 | 2.299 |
| 9fu7_NI_multi_hetero_d7.00_cluster1_altlocLIGA.xyz | 2.578 | | 2.805 | 2.11 | 2.17 | 2.313 |
| 9fu7_NI_multi_hetero_d7.00_cluster1_altlocLIGB.xyz | 2.578 | | 2.805 | 2.11 | 2.17 | 2.313 |
| 9fop_NI_multi_hetero_d7.00_cluster1.xyz | 2.536 | | 2.807 | 2.11 | 2.181 | 2.297 |
| 9fua_NI_multi_hetero_d7.00_cluster1.xyz | 2.582 | | 2.807 | 2.111 | 2.187 | 2.297 |
| 1mjg_NI_multi_hetero_d7.00_cluster4.xyz | 2.684 | | 2.809 | 2.23 | 2.267 | 2.3 |
| 9fox_NI_multi_hetero_d7.00_cluster2.xyz | 2.527 | | 2.81 | 2.097 | 2.18 | 2.289 |
| 4udx_NI_multi_hetero_d7.00_cluster1_altlocLIGA.xyz | 2.079 | | 2.813 | 2.112 | 2.2 | 2.331 |
| 9fr1_NI_multi_hetero_d7.00_cluster2.xyz | 2.511 | | 2.813 | 2.109 | 2.177 | 2.293 |
| 9fnc_NI_multi_hetero_d7.00_cluster2.xyz | 2.522 | | 2.813 | 2.109 | 2.185 | 2.3 |
| 9fr1_NI_multi_hetero_d7.00_cluster1_altlocLIGA.xyz | 2.644 | | 2.813 | 2.098 | 2.176 | 2.297 |
| 9fr1_NI_multi_hetero_d7.00_cluster1_altlocLIGB.xyz | 2.644 | | 2.813 | 2.098 | 2.176 | 2.297 |
| 9fo4_NI_multi_hetero_d7.00_cluster2.xyz | 2.665 | | 2.813 | 2.11 | 2.182 | 2.293 |
| 4udx_NI_multi_hetero_d7.00_cluster1_altlocLIGB.xyz | 2.829 | | 2.813 | 2.2 | 2.331 | 2.918 |
| 9fu4_NI_multi_hetero_d7.00_cluster1_altlocLIGA.xyz | 2.63 | | 2.814 | 2.036 | 2.182 | 2.298 |
| 9fu4_NI_multi_hetero_d7.00_cluster1_altlocLIGB.xyz | 2.63 | | 2.814 | 2.036 | 2.182 | 2.298 |
| 9fuc_NI_multi_hetero_d7.00_cluster1.xyz | 2.471 | | 2.815 | 2.11 | 2.166 | 2.296 |
| 9fop_NI_multi_hetero_d7.00_cluster2.xyz | 2.673 | | 2.815 | 2.11 | 2.177 | 2.297 |
| 3i39_NI_multi_hetero_d7.00_cluster1_altlocLIGA.xyz | 2.558 | 3.007 | 2.817 | 2.079 | 2.17 | 2.264 |
| 3i39_NI_multi_hetero_d7.00_cluster1_altlocLIGB.xyz | 2.558 | 3.007 | 2.817 | 2.079 | 2.17 | 2.264 |
| 9fnj_NI_multi_hetero_d7.00_cluster2.xyz | 2.609 | | 2.82 | 2.11 | 2.19 | 2.303 |
| 1mjg_NI_multi_hetero_d7.00_cluster3.xyz | 2.606 | | 2.822 | 2.136 | 2.282 | 2.288 |
| 9fox_NI_multi_hetero_d7.00_cluster1.xyz | 2.528 | | 2.823 | 2.109 | 2.184 | 2.288 |
| 5fle_NI_multi_hetero_d7.00_cluster1_altlocLIGA.xyz | 2.67 | | 2.823 | 2.118 | 2.201 | 2.292 |
| 5fle_NI_multi_hetero_d7.00_cluster1_altlocLIGB.xyz | 2.67 | | 2.823 | 2.118 | 2.201 | 2.292 |
| 9fo4_NI_multi_hetero_d7.00_cluster1.xyz | 2.614 | | 2.825 | 2.109 | 2.183 | 2.299 |
| 9fph_NI_multi_hetero_d7.00_cluster1_altlocLIGA.xyz | 2.805 | | 2.825 | 2.142 | 2.204 | 2.312 |
| 3i04_NI_multi_hetero_d7.00_cluster1.xyz | 3.124 | | 2.836 | 2.142 | 2.325 | 2.334 |
| 9fu3_NI_multi_hetero_d7.00_cluster1.xyz | 2.565 | | 2.842 | 2.11 | 2.173 | 2.3 |
| 4udy_NI_multi_hetero_d7.00_cluster1_altlocLIGA.xyz | 2.089 | | 2.849 | 2.102 | 2.211 | 2.312 |
| 4udy_NI_multi_hetero_d7.00_cluster1_altlocLIGB.xyz | 2.904 | | 2.849 | 2.102 | 2.211 | 2.312 |
| 3i04_NI_multi_hetero_d7.00_cluster3.xyz | 3.121 | | 2.851 | 2.102 | 2.213 | 2.442 |
| 6x5k_NI_multi_hetero_d7.00_cluster3_altlocLIGA.xyz | 2.931 | | 2.852 | 2.147 | 2.346 | 2.347 |
| 9fpg_NI_multi_hetero_d7.00_cluster1_altlocLIGA.xyz | 2.867 | | 2.853 | 2.048 | 2.174 | 2.27 |
| 8x9f_NI_multi_hetero_d7.00_cluster1.xyz | 2.644 | | 2.854 | 1.981 | 2.184 | 2.465 |
| 2yiv_NI_multi_hetero_d7.00_cluster1_altlocLIGB.xyz | 2.193 | | 2.855 | 2.092 | 2.166 | 2.321 |
| 2yiv_NI_multi_hetero_d7.00_cluster1_altlocLIGA.xyz | 3.105 | | 2.855 | 2.092 | 2.166 | 2.321 |
| 7zy1_NI_multi_hetero_d7.00_cluster1_altlocNiA.xyz | 2.375 | | 2.894 | 2.085 | 2.208 | 2.356 |
| 9gyb_NI_multi_hetero_d7.00_cluster3.xyz | 2.625 | | 2.902 | 1.973 | 2.316 | 2.321 |
| 8x9d_NI_multi_hetero_d7.00_cluster1_altlocLIGA.xyz | 2.84 | | 2.906 | 1.975 | 2.182 | 2.438 |
| 8x9d_NI_multi_hetero_d7.00_cluster1_altlocLIGB.xyz | 2.84 | | 2.906 | 1.985 | 2.182 | 2.438 |
| 8x9h_NI_multi_hetero_d7.00_cluster1.xyz | 2.827 | | 2.922 | 1.98 | 2.194 | 2.441 |
| 9iyr_NI_multi_hetero_d7.00_cluster1_altlocLIGA.xyz | 2.925 | | 2.948 | 1.975 | 2.223 | 2.511 |
| 9iyr_NI_multi_hetero_d7.00_cluster1_altlocLIGB.xyz | 2.925 | | 2.948 | 1.989 | 2.223 | 2.511 |
| 9fzy_NI_multi_hetero_d7.00_cluster2.xyz | 2.502 | | 2.955 | 2.068 | 2.298 | 2.39 |
| 9fzz_NI_multi_hetero_d7.00_cluster2.xyz | 2.502 | | 2.955 | 2.085 | 2.298 | 2.39 |
| 9g00_NI_multi_hetero_d7.00_cluster2.xyz | 2.502 | | 2.955 | 2.085 | 2.298 | 2.39 |
| 9g01_NI_multi_hetero_d7.00_cluster1.xyz | 2.502 | | 2.955 | 2.084 | 2.298 | 2.39 |
| 9g02_NI_multi_hetero_d7.00_cluster2.xyz | 2.502 | | 2.955 | 2.085 | 2.298 | 2.39 |
| 9g03_NI_multi_hetero_d7.00_cluster1.xyz | 2.502 | | 2.955 | 2.069 | 2.298 | 2.39 |
| 9iym_NI_multi_hetero_d7.00_cluster1_altlocLIGA.xyz | 2.762 | | 2.957 | 1.976 | 2.195 | 2.467 |
| 9iym_NI_multi_hetero_d7.00_cluster1_altlocLIGB.xyz | 2.762 | | 2.957 | 1.993 | 2.195 | 2.467 |
| 9g03_NI_multi_hetero_d7.00_cluster2.xyz | 2.503 | | 2.959 | 2.082 | 2.297 | 2.399 |
| 9fzy_NI_multi_hetero_d7.00_cluster3.xyz | 2.504 | | 2.959 | 2.059 | 2.297 | 2.399 |
| 9fzz_NI_multi_hetero_d7.00_cluster3.xyz | 2.504 | | 2.959 | 2.097 | 2.297 | 2.399 |
| 9g00_NI_multi_hetero_d7.00_cluster3.xyz | 2.504 | | 2.959 | 2.096 | 2.297 | 2.399 |
| 9g01_NI_multi_hetero_d7.00_cluster2.xyz | 2.504 | | 2.959 | 2.097 | 2.297 | 2.399 |
| 9g02_NI_multi_hetero_d7.00_cluster3.xyz | 2.504 | | 2.959 | 2.097 | 2.297 | 2.399 |
| 6yu9_NI_multi_hetero_d7.00_cluster3.xyz | 2.675 | | 2.976 | 2.247 | 2.251 | 2.541 |
| 9iys_NI_multi_hetero_d7.00_cluster1_altlocLIGA.xyz | 2.895 | | 2.978 | 1.976 | 2.22 | 2.623 |
| 9iys_NI_multi_hetero_d7.00_cluster1_altlocLIGB.xyz | 2.895 | | 2.978 | 1.991 | 2.22 | 2.623 |
| 9iyo_NI_multi_hetero_d7.00_cluster1_altlocLIGA.xyz | 2.876 | | 2.988 | 1.981 | 2.19 | 2.455 |
| 9iyo_NI_multi_hetero_d7.00_cluster1_altlocLIGB.xyz | 2.876 | | 2.988 | 1.982 | 2.19 | 2.455 |
| 9iyn_NI_multi_hetero_d7.00_cluster1_altlocLIGA.xyz | 2.765 | | 3.1 | 1.97 | 2.208 | 2.578 |
| 9iyn_NI_multi_hetero_d7.00_cluster1_altlocLIGB.xyz | 2.765 | | 3.1 | 1.988 | 2.208 | 2.578 |
| 2z8y_NI_multi_hetero_d7.00_cluster1.xyz | 2.935 | | 3.139 | 2.17 | 2.352 | 2.412 |
| 9iyt_NI_multi_hetero_d7.00_cluster1_altlocLIGA.xyz | 2.723 | | 3.141 | 1.976 | 2.213 | 2.671 |
| 9iyt_NI_multi_hetero_d7.00_cluster1_altlocLIGB.xyz | 2.723 | | 3.141 | 1.984 | 2.213 | 2.671 |
| 9iyl_NI_multi_hetero_d7.00_cluster1_altlocLIGA.xyz | 3.088 | | 3.178 | 1.978 | 2.238 | 2.567 |
| 9iyl_NI_multi_hetero_d7.00_cluster1_altlocLIGB.xyz | 3.088 | | 3.178 | 1.991 | 2.238 | 2.567 |
| 2z8y_NI_multi_hetero_d7.00_cluster4.xyz | 3.103 | | 3.199 | 2.096 | 2.381 | 2.404 |
| 9iyu_NI_multi_hetero_d7.00_cluster1_altlocLIGA.xyz | 2.915 | | 3.237 | 1.988 | 2.219 | 2.708 |
| 9iyu_NI_multi_hetero_d7.00_cluster1_altlocLIGC.xyz | 2.915 | | 3.237 | 1.978 | 2.219 | 2.708 |
| 2z8y_NI_multi_hetero_d7.00_cluster2.xyz | 3.073 | | 3.238 | 2.167 | 2.363 | 2.416 |
| 2z8y_NI_multi_hetero_d7.00_cluster3.xyz | 3.152 | | 3.261 | 1.785 | 2.325 | 2.392 |
| 8cmw_NI_multi_hetero_d7.00_cluster1_altlocLIGA.xyz | 2.363 | | 3.272 | 1.958 | 2.285 | 2.492 |
| 8cmw_NI_multi_hetero_d7.00_cluster1_altlocLIGB.xyz | 2.363 | | 3.272 | 2.285 | 2.492 | 2.698 |
| 8cmw_NI_multi_hetero_d7.00_cluster1_altlocLIGC.xyz | 2.363 | | 3.272 | 2.285 | 2.492 | 2.719 |
| 9g7i_NI_multi_hetero_d7.00_cluster3.xyz | 2.633 | | 3.389 | 2.253 | 2.501 | 2.862 |
| 8cja_NI_multi_hetero_d7.00_cluster1_altlocLIGA.xyz | 2.457 | | 3.426 | 1.897 | 2.339 | 2.618 |
| 8cja_NI_multi_hetero_d7.00_cluster1_altlocLIGB.xyz | 2.457 | | 3.426 | 2.339 | 2.618 | 2.898 |
| 8cja_NI_multi_hetero_d7.00_cluster1_altlocLIGC.xyz | 2.457 | | 3.426 | 2.339 | 2.591 | 2.618 |
| 6ytt_NI_multi_hetero_d7.00_cluster2.xyz | 2.577 | 4.148 | 3.426 | 2.291 | 2.695 | 2.976 |
| 9g7i_NI_multi_hetero_d7.00_cluster2.xyz | 2.577 | | 3.426 | 2.253 | 2.291 | 2.695 |
| 6ytt_NI_multi_hetero_d7.00_cluster3.xyz | 2.511 | | 3.448 | 2.287 | 2.58 | 2.975 |
| 8cjc_NI_multi_hetero_d7.00_cluster1_altlocLIGA.xyz | 2.391 | | 3.574 | 1.911 | 2.395 | 2.639 |
| 8cjc_NI_multi_hetero_d7.00_cluster1_altlocLIGB.xyz | 2.391 | | 3.574 | 2.395 | 2.527 | 2.639 |

Table S2. EPR simulation parameters

| **B Cluster (-430 mV)** | g1 | g2 | g3 | g1 strain | g2 strain | g3 strain | Proportion |
|---|---|---|---|---|---|---|---|
| B Cluster | 1.892 | 1.934 | 2.036 | 0.025 | 0.020 | 0.025 | 0.911 |
| C cluster | 1.740 | 1.830 | 1.990 | 0.053 | 0.054 | 0.022 | 0.089 |
| **Cred1 (-370 mV)** | g1 | g2 | g3 | g1 strain | g2 strain | g3 strain | Proportion |
| B Cluster | 1.893 | 1.932 | 2.036 | 0.028 | 0.014 | 0.025 | 0.345 |
| C cluster Conformation 1 | 1.635 | 1.849 | 1.968 | 0.045 | 0.073 | 0.021 | 0.484 |
| C cluster Conformation 2 | 1.735 | 1.813 | 2.002 | 0.030 | 0.038 | 0.022 | 0.164 |
| Benzyl Viologen | 2.003 | – | – | 0.015 | – | – | 0.008 |
| **Cred2 (-540 mV)** | g1 | g2 | g3 | g1 strain | g2 strain | g3 strain | Proportion |
| B Cluster | 1.891 | 1.931 | 2.034 | 0.026 | 0.016 | 0.022 | 0.796 |
| C cluster | 1.727 | 1.811 | 1.969 | 0.044 | 0.046 | 0.020 | 0.122 |
| D cluster | 1.862 | 1.951 | 2.006 | 0.031 | 0.028 | 0.018 | 0.082 |
| **Cred1+CN** | g1 | g2 | g3 | g1 strain | g2 strain | g3 strain | Proportion |
| B Cluster | 1.895 | 1.933 | 2.039 | 0.019 | 0.022 | 0.020 | 0.371 |
| C cluster | 1.849 | 1.883 | 2.011 | 0.054 | 0.038 | 0.025 | 0.617 |
| Benzyl Viologen | 2.003 | – | – | 0.013 | – | – | 0.012 |
| **Cred2+OCN** | g1 | g2 | g3 | g1 strain | g2 strain | g3 strain | Proportion |
| B Cluster | 1.893 | 1.931 | 2.036 | 0.029 | 0.019 | 0.024 | 0.709 |
| C cluster | 1.733 | 1.851 | 1.970 | 0.044 | 0.067 | 0.025 | 0.235 |
| D cluster | 1.869 | 1.936 | 2.003 | 0.025 | 0.054 | 0.015 | 0.057 |

Table S3. Psuedo-Voigt Peak Deconvolution of Fe K-edge Datasets

| Species | edge (μE = 0.5) | Pre-edg fit centroid | Peak 1 amp | Peak 2 amp | PE amplitude | peak 1 energy | peak 2 energy | delta | $\chi^2$ |
|---|---|---|---|---|---|---|---|---|---|
| Cox | 7118.73 | 7112.69 | 0.200 | 0.079 | 0.278 | 7112.39 | 7113.45 | 1.06 | 0.0001 |
| Cred1 | 7118.55 | 7112.53 | 0.183 | 0.063 | 0.246 | 7112.29 | 7113.23 | 0.94 | 0.0003 |
| Cred2 | 7118.31 | 7112.21 | 0.210 | – | 0.210 | 7112.21 | – | – | 0.0005 |
| Cred1+CN | 7118.51 | 7112.50 | 0.189 | 0.053 | 0.243 | 7112.28 | 7113.25 | 0.97 | 0.0003 |
| Cred2+OCN | 7118.42 | 7112.41 | 0.157 | 0.047 | 0.204 | 7112.22 | 7113.06 | 0.84 | 0.0004 |
| K4[Fe4S4]0 | 7117.4 | 7111.88 | 0.060 | 0.087 | 0.147 | 7111.65 | 7112.05 | 0.40 | 0.0005 |
| K3[Fe4S4]+1 | 7117.91 | 7112.08 | 0.114 | 0.077 | 0.191 | 7111.80 | 7112.50 | 0.70 | 0.0002 |
| K2[Fe4S4]+2 | 7118.24 | 7112.35 | 0.162 | 0.080 | 0.243 | 7112.04 | 7112.98 | 0.94 | 0.0000 |
| C Clsuter - Cox | 7118.4 | 7112.82 | 0.234 | 0.086 | 0.320 | 7112.53 | 7113.60 | 1.07 | 0.0002 |
| C Clsuter - Cred1 | 7117.98 | 7112.51 | 0.208 | 0.044 | 0.252 | 7112.34 | 7113.29 | 0.95 | 0.0004 |
| C Clsuter - Cred2 | 7117.63 | 7112.22 | 0.209 | 0.003 | 0.212 | 7112.20 | 7113.42 | 1.22 | 0.0005 |

| Table S4. Psuedo-Voigt Peak Deconvolution of Fe HERFD Pre-Edge | | | | | |
|---|---|---|---|---|---|
| Iron (HERFD Si311) | edge (μE = 0.5) | Pseudo-Voigt Centroid | Pseudo-Voigt 1 Amplitude | Pseudo-Voigt 2 Amplitude | Total Amplitude |
| Cox | 7118.47 | 7112.34 | 0.340 | 0.050 | 0.391 |
| Cred1 | 7118.3 | 7112.18 | 0.266 | 0.071 | 0.337 |
| Cred2 | 7117.94 | 7112.04 | 0.248 | 0.059 | 0.307 |

Table S5. Assesment of existing substrate-free CODH-II crystal structures. Values in bold are the maximum RSZD+/- score for the selected side chain, or cluster*. Values in parentheses are the atomic occupancy. Values in curly brackets are the atomic B-factor.

| Atom | 3b51 (Cred2) | 3b53 (Cred1) | 1su7 (Cred1) | 1suf (Inactive state) | 1su8 (Cred2) | 9FPG (Cred1) | Atom Numbering |
|---|---|---|---|---|---|---|---|
| S1 | (0.8) {7.63} | (0.8) {12.55} | (1) {13.44} | (1) {11.25} | (1) {11.98} | (0.79) {18.3} | |
| S2 | (0.8) {8.38} | (0.8) {13.06} | (0.95) {12.64} | (1) {13.31} | (0.95) {12.03} | (0.75) {17.28} | |
| S3 | (0.8) {8.61} | (0.8) {11.81} | (0.98) {15.51} | (1) {11.88} | (0.98) {14.05} | (0.78) {17.73} | |
| S4 | (0.8) {7.99} | (0.8) {11.46} | (0.98) {12.27} | (1) {10.75} | 9 | (0.79) {17.51} | |
| S5 (Ni-Feu bridge) | (x) {x} | (x) {x} | (0.83) {15.69} | (x) {x} | (0.83) {14.31} | (x) {x} | |
| O | x (x) {x} | x (x) {x} | 11.1 (1) {34.8} | 2.7 (1) {25.32} | 6.5 (1) {49.27} | x (0.67) {17.93} | |
| O | x (x) {x} | x (x) {x} | x (x) {x} | -2.7 (1) {27.73} | x (x) {x} | x (x) {x} | |
| O (Feu) | **-1.3** (0.6) {10.24} | **2.9** (0.6) {10.34} | **x** (x) {x} | **-3.1** (1) {21.38} | **x** (x) {x} | **2.3** (0.67) {17.93} | |
| Feu i/o | **-5.7** (0.6/0.3) {10.76/8.17} | **-3.6** (0.6/0.3) {12.63/12.52} | **x** (0.9) {12.6} | **x** (0.6/0.3) {4.28/7.64} | **x** (0.8/0.1) {11.12/8.96} | **x** (0.62/0.23) {15.84/14.75} | |
| Fe2 | (0.8) {8.35} | (0.8) {12.55} | (0.98) {9.98} | (1) {0.3} | (0.98) {9.61} | (0.8) {17.19} | |
| Fe3 | (0.8) {9.3} | (0.8) {12.97} | (0.7/0.3) {10.08/26.80} | (1) {0.48} | (0.85/0.1) {10.39/10.08} | (0.83) {18.36} | |
| Fe4 | (0.8) {7.86} | (0.8) {11.71} | (0.95) {10.43} | (1) {0.08} | (0.95) {10.11} | (0.75) {16.03} | |
| Ni | (0.6) {10.77} | (0.6) {13.64} | (0.93) {12.45} | (0.6) {1.58} | (0.93) {11.35} | (0.66) {18.15} | |
| Cys526 | **3.7** (1) {9.8} | **-5** (1) {11.45} | **-18.2** (1) {14.85} | **2.1** (1) {10.93} | **-11.7** (1) {12.25} | **-7.1** (0.85/0.15) {17.28/17.63} | |
| Cys446 | **1.5** (1) {7.84} | **1.8** (1) {10.63} | **-0.7** (1) {9.44} | **6.2** (1) {10.36} | **-0.6** (1) {9.36} | **-1.6** (1) {16.82} | |
| Cys476 | **3.1** (1) {13.02} | **2.6** (1) {17.04} | **-5.3** (1) {13.21} | **4.3** (1) {11.85} | **-2.7** (1) {12.58} | **0.7** (0.78/0.22) {18.09/17.17} | |
| Cys295 a/b | **0.6** (0.7/0.3) {10.68/5.57} | **2.1** (0.7/0.3) {13.19/9.82} | **-2.2** (1) {11.71} | **3.4** (1) {13.81} | **2.2** (1) {11.99} | **1.2** (0.6/0.4) {14.70/15.69} | |
| Cys333 | **-0.7** (1) {7.75} | **1.5** (1) {11.56} | **-6.7** (1) {10.11} | **5.3** (1) {10.08} | **-2.3** (1) {10.54} | **-6.3** (1) {16.77} | |
| **WCC*** | **4.2** | **-4.3** | **x** | **x** | **x** | **-7.9** | |
| **NFS**** | **x** | **x** | **-99.9, 8.4** | **-99.9, 8.9** | **-99.9, 1.5** | **x** | |

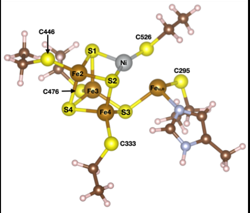

* In the case of protein ligands, such as the C cluster, the RSZD is scored based on how the ligand is described in the pdb structure file. In 1SUF, 1SU7 and 1SU8, the whole C cluster (NiFe4S4) is descibed in the NFS ligand, whereas Feu was split out as a separate ligand in 3B51, 3B53 and 9FPG allowing it to be scored seperately from the remaining (NiFe3S4). A RSZD score <-3σ and >3σ are unlikely to occur as a result of chance random errors and therefore, an advisable rejection limit. ** Due to the large value of RSDZ-, both the RSDZ+ and RSDZ- are shown.

| Table S6. C cluster Ni-(S,L,Fe) distances for the available substrate-free CODH crystal structures (Å) | | | | | | | |
|---|---|---|---|---|---|---|---|
| | 3b51.pdb | 3b53.pdb | 1suf.pdb | 1su7.pdb | 1su8.pdb | 9FPG | Atom Numbering |
| Species | -600 MV STATE (Cred2) | -320 MV STATE (Cred1) | INACTIVE STATE | educed (Cred1) N | CODHIIch incubated with N2_Dithionite | Ti(III) citrate Cred1 pH 6 | 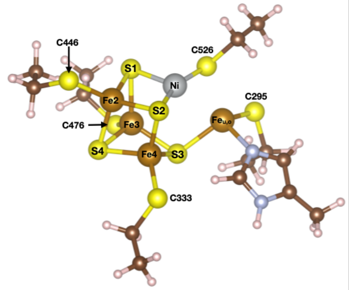 |
| Cys526 | 2.17 | 2.17 | 2.14 | 2.32 | 2.26 | 2.05 | |
| S1 | 2.16 | 2.14 | 2.44 | 2.3 | 2.31 | 2.27 | |
| S2 | 2.03 | 2.07 | 2.18 | 2.24 | 2.25 | 2.17 | |
| Avg Ni-S | 2.12 | 2.13 | 2.25 | 2.29 | 2.27 | 2.16 | |
| S3 | 3.53 | 3.47 | 4.06 | 3.66 | 3.75 | 3.75 | |
| S4 | 4.38 | 4.36 | 4.78 | 4.59 | 4.62 | 4.59 | |
| S (bridging) | - | - | - | 2.21 | 2.28 | - | |
| O | - | - | 2.84 | 2.29 | 2.28 | 2.24 (bridging) | |
| O | - | - | 2.84 | - | - | - | |
| O (Feu) | 2.7 | 2.72 | 2.95 | - | - | 2.05 (bridging) | |
| Feu (out) | 2.87 | 2.85 | 3.11 | 2.84 | 2.82 | 2.87 | |
| Fe2 | 2.62 | 2.65 | 2.79 | 2.84 | 2.88 | 2.85 | |
| Fe3 | 3.42 | 3.37 | 3.82 | 3.67/5.04 | 3.47/3.62 | 3.53 | |
| Fe4 | 3.18 | 3.09 | 3.6 | 3.34 | 3.34 | 3.3 | |

| Table 7. EXAFS least-squares fitting parameters for fits presented in Figure S5. | | | | | | | | |
|---|---|---|---|---|---|---|---|---|
| Fit | Path | Coordination | $S_0^2$ | R(Å)[a] | $\sigma^2$ (Å$^2$)[b] | ΔE | K range | F-factor |
| Cox-c | S | 3 | 0.9 | 2.22 | 477 | -0.4 | 2-16 | 0.36 |
| Cox-d | S | 2 | 0.9 | 2.20 | 207 | 1.5 | 2-16 | 0.33 |
| | S | 1 | | 2.30 | 154 | | | |
| | Fe | 1 | | 2.68 | 1103 | | | |
| | Fe | 1 | | 2.91 | 1152 | | | |
| Cox-e | S | 3 | 0.9 | 2.23 | 649 | 4.3 | 2-16 | 0.33 |
| | O | 1 | | 2.09 | 134 | | | |
| | Fe | 1 | | 2.70 | 1097 | | | |
| | Fe | 1 | | 2.93 | 1048 | | | |
| Cred1-f | S | 3 | 0.9 | 2.20 | 326 | -4.7 | 2-16 | 0.35 |
| | O | 2 | | 2.41 | 199 | | | |
| Cred1-g | S | 3 | 0.9 | 2.23 | 301 | 1.2 | 2-16 | 0.36 |
| | Fe | 1 | | 2.71 | 714 | | | |
| | Fe | 1 | | 2.92 | 789 | | | |
| Cred1-h | S | 3 | 0.9 | 2.21 | 303 | -3.0 | 2-16 | 0.32 |
| | O | 1 | | 2.43 | 202 | | | |
| | Fe | 1 | | 2.67 | 634 | | | |
| | Fe | 1 | | 2.88 | 938 | | | |
| Cred1+CN-h | S | 3 | 0.9 | 2.21 | 492 | 1.1 | 2-16 | 0.29 |
| | C | 1 | | 1.84 | 13 | | | |
| | Fe | 1 | | 2.68 | 583 | | | |
| | Fe | 2 | | 2.83 | 453 | | | |

[a]The estimated standard deviations in interatomic distance is ± 0.02 Å. [b]The $\sigma^2$ values have been multiplied by $10^5$. a / indicates the value of the path was tied to the one above it.

| Table S8. Ni-Fe distances obtained from Least-squares fits to the Ni K-edge EXAFS (Å) | | | | | |
|---|---|---|---|---|---|
| Sample | Fe(u) | Fe(2) | Fe | Fe | Fe2-Feu (Å) |
| Cox | 2.68 | 2.91 | – | – | 0.23 |
| Cred1 | 2.67 | 2.88 | – | – | 0.21 |
| Cred2 | 2.57 | 2.75 | – | – | 0.18 |
| Cred1+CN | 2.69 | 2.83 | 2.83 | 3.19 | 0.14 |
| Cred2+OCN | 2.75 | 2.92 | – | – | 0.17 |

Table S9. Crystallographic Ni-Fe distances from CODH-II obtained from Basak et al. 2025

| Distance to Ni | Feu,o | Feu,i | Ni occ | Feu,o occ | Feu,I occ |
|---|---|---|---|---|---|
| 9FPF | 2.88 | 2.41 | 0.73 | 0.53 | 0.35 |
| 9fpo | 2.87 | 2.26 | 0.24 | 0.55 | 0.3 |
| 9fpn | – | 2.6 | 0.78 | – | 0.78 |
| 9fpl | 2.76 | 2.18 | 0.5 | 0.21 | 0.56 |
| 9fpk | 2.83 | 2.29 | 0.34 | 0.57 | 0.25 |
| 9fpj | 2.84 | 2.23 | 0.6 | 0.31 | 0.6 |
| 9fpi | 2.8 | 2.07 | 0.31 | 0.5 | 0.37 |
| 9fph | 2.8 | 2.3 | 0.66 | 0.2 | 0.63 |
| 9fpg | 2.87 | 2.38 | 0.66 | 0.62 | 0.23 |

| Table S10. EPR data acquisition parameters | | | | | | |
|---|---|---|---|---|---|---|
| Sample | Cred1 (-350 mW) | Cred1 (-350 mW) | –430 mV | Cred2 (-540 mV) | Cred1+CN | Cred2+OCN |
| Mircowave Frequencey (GHz) | 9.37 | 9.38 | 9.38 | 9.37 | 9.38 | 9.38 |
| Mircowave power (mW) | 20.00 | 46.00 | 20.00 | 2.00 | 46.00 | 54.00 |
| Modulation Amplitude (G) | 10 | 8 | 10 | 10 | 8 | 8 |
| Conversion time (ms) | 163.84 | 655.36 | 163.84 | 163.84 | 655.36 | 655.36 |
| RF Attenuation (dB) | 10 | 6 | 12 | 20 | 12 | 4 |
| Time constant (ms) | 81.92 | 81.92 | 81.92 | 81.92 | 81.92 | 81.92 |
| Reciever Gain | 5023.77 | 1415.89 | 5023.77 | 5023.77 | 1415.89 | 1415.89 |

| Table S11. Summary values from the PDB survey used to construct Figure 2 | | | | | |
|---|---|---|---|---|---|
| | Feu | Fe2 | S1 | S2 | S3 |
| Median | 2.64 | 2.81 | 2.11 | 2.22 | 2.34 |
| Mean | 2.70 | 2.86 | 2.11 | 2.25 | 2.40 |
| Stdev | 0.25 | 0.22 | 0.11 | 0.11 | 0.17 |

| Table S12. RSZD Scoring of all CODH C-clsuters available in the PDB | | |
|---|---|---|
| **PDB** | **C Cluster RSDZ +/-** | **C Cluster RSDZ Score** |
| 1SU7 | -99.9 | 99.9 |
| 1SU8 | -99.9 | 99.9 |
| 1SUF | -99.9 | 99.9 |
| 7XDP | -32.3 | 32.3 |
| 7XDN | -29.2 | 29.2 |
| 7XDM | -28.8 | 28.8 |
| 8X9D | -21.3 | 21.3 |
| 2YIV | -17.6 | 17.6 |
| 6OND | -17.1 | 17.1 |
| 6B6X | -16.5 | 16.5 |
| 7B97 | -10.9 | 10.9 |
| 8X9E | -10.7 | 10.7 |
| 7ZX5 | -9.8 | 9.8 |
| 9FPJ | -9.3 | 9.3 |
| 9FPK | -8.9 | 8.9 |
| 1SU6 | -8.6 | 8.6 |
| 9FPN | -8 | 8 |
| 9FPG | -7.9 | 7.9 |
| 7ZXL | -7.6 | 7.6 |
| 7ZXJ | -7 | 7 |
| 9FPF | -6.9 | 6.9 |
| 3I39 | -6.6 | 6.6 |
| 8X9H | -5.2 | 5.2 |
| 7B7Q | -4.9 | 4.9 |
| 6DC2 | -4.7 | 4.7 |
| 7ZXC | -4.7 | 4.7 |
| 3B53 | -4.3 | 4.3 |
| 3B51 | -4.2 | 4.2 |
| 3B52 | -3.4 | 3.4 |
| 8OMX | -3.4 | 3.4 |
| 8X9F | -3.1 | 3.1 |
| 8X9G | -2.9 | 2.9 |
| 7XY1 | -2.3 | 2.3 |
| 6B6V | -1.5 | 1.5 |
| 5FLE | -1.4 | 1.4 |
| 6T7J | 1.2 | 1.2 |
| 7B9A | 1.8 | 1.8 |
| 9IYV | 2.8 | 2.8 |
| 6ELQ | 2.9 | 2.9 |
| 9FPL | 3.3 | 3.3 |
| 7ZX6 | 3.7 | 3.7 |
| 7ERR | 3.7 | 3.7 |
| 6B6W | 4.3 | 4.3 |
| 9IYO | 4.3 | 4.3 |
| 9IYU | 4.5 | 4.5 |
| 4UDX | 5.6 | 5.6 |
| 9IYN | 5.6 | 5.6 |
| 4UDY | 6.6 | 6.6 |
| 9IYT | 6.6 | 6.6 |
| 6YU9 | 6.7 | 6.7 |
| 9FPI | 6.9 | 6.9 |
| 7B7T | 7.3 | 7.3 |
| 9IYM | 7.7 | 7.7 |
| 7TSJ | 8.4 | 8.4 |
| 7B95 | 8.9 | 8.9 |
| 6ONS | 9.4 | 9.4 |
| 9iYR | 9.5 | 9.5 |
| 9FPO | 10 | 10 |
| 9IYS | 12.2 | 12.2 |
| 9FPH | 12.5 | 12.5 |
| 9IYL | 14.2 | 14.2 |
| 6B6Y | 1.8, 2.9 | 1.8, 2.9 |
| 9GYB | 5.4, 3.8, 3.8, 6.1, 1.1,2.7 | 5.4, 3.8, 3.8, 6.1, 1.1,2.7 |
